# Vesicular stomatitis virus transcription is inhibited by TRIM69 in the interferon-induced antiviral state

**DOI:** 10.1101/683292

**Authors:** Tonya Kueck, Louis-Marie Bloyet, Elena Cassella, Trinity Zang, Fabian Schmidt, Vesna Brusic, Gergely Tekes, Owen Pornillos, Sean P. J. Whelan, Paul D. Bieniasz

## Abstract

Interferons (IFNs) induce the expression of many interferon stimulated genes (ISGs), many of which are responsible for the cellular ‘antiviral state’ in which the replication of numerous viruses is blocked. How the majority of individual ISGs inhibit the replication of particular viruses is unknown. We conducted a loss-of-function screen to identify genes required for the activity of IFN*α* against vesicular stomatitis virus, Indiana serotype (VSV_IND_), a prototype negative strand RNA virus. Our screen revealed that TRIM69, a member of tripartite motif family of proteins, is a VSV_IND_ inhibitor. TRIM69 potently inhibited VSV_IND_ replication through a previously undescribed transcriptional inhibition mechanism. Specifically, TRIM69 physically associates with the VSV_IND_ phosphoprotein (P), requiring a specific peptide target sequence encoded therein. P is a cofactor for the viral polymerase, and is required for viral RNA synthesis as well as the assembly of replication compartments. By targeting P, TRIM69 inhibits pioneer transcription of the incoming virion-associated minus strand RNA, thereby preventing the synthesis of viral mRNAs, and consequently impedes all downstream events in the VSV_IND_ replication cycle. Unlike some TRIM proteins, TRIM69 does not inhibit viral replication by inducing degradation of target viral proteins. Rather, higher-order TRIM69 multimerization is required for its antiviral activity, suggesting that TRIM69 functions by sequestration or anatomical disruption of the viral machinery required for VSV_IND_ RNA synthesis.

**SIGNIFICANCE STATEMENT:** Interferons are important antiviral cytokines that work by inducing hundreds of host genes whose products inhibit replication of many viruses. While the antiviral activity of interferon has long been known, the identities and mechanisms of action of most interferon-induced antiviral proteins remain to be discovered. We identified gene products that are important for the antiviral activity of interferon against vesicular stomatitis virus (VSV) a model virus that whose genome consists a single RNA molecule with negative sense polarity. We found that a particular antiviral protein, TRIM69, functions by a previously undescribed molecular mechanism. Specifically, TRIM69 interacts with, and inhibits the function, of a particular phosphoprotein (P) component the viral transcription machinery, preventing the synthesis of viral messenger RNAs.

---

Infection of vertebrate animal cells by many viruses triggers innate immune responses, among which the induction of type-I interferons (IFNα/β) is especially important. IFNs induce the expression of hundreds of interferon stimulated genes (ISGs) that have a plethora of downstream effects in a wide range of cells (1). In particular, the products of many ISGs contribute to the establishment of the so called ‘antiviral state’, in which the replication of many viruses is blocked (2, 3). Some ISGs have been shown to inhibit specific processes in viral replication, or induce the destruction or depletion of viral proteins or RNA. However, the function of the majority of individual ISGs, and precisely how they inhibit replication of particular viruses, remains unknown (4).

Vesicular stomatitis virus (VSV), a prototypic member of the order *Mononegavirales*, and an important animal pathogen, is highly sensitive to inhibition by type-I IFN. Like many RNA viruses, VSV replication occurs within specialized compartments in the cytoplasm of infected cells, (5, 6). Compartmentalization may help to shield viral components from detection by cytosolic sensors or antiviral proteins that might otherwise increase IFN production or directly interfere with viral replication. For rhabdoviruses such as VSV and rabies virus, and other cytoplasmically replicating negative strand RNA viruses, replication compartments are not circumscribed by a membrane (6–9). Instead, replication components form inclusions that manifest features characteristic of phase-separated liquid compartments such as P-bodies and nucleoli (10, 11). Three VSV proteins, namely the nucleoprotein (N) which coats the viral RNA, the large protein (L) that possesses all the viral enzymes necessary for transcription and the phosphoprotein (P) that binds to both N and L to stimulate RNA synthesis, are necessary and sufficient for the assembly of phase-separated replication compartments (10). While not required for compartment formation, negative strand viral RNA is located, replicated and transcribed within these compartments once infection is established (6). However, the initial ‘pioneer’ round of transcription, in which viral mRNAs are transcribed from a single incoming negative strand viral genome, employs N, P and L proteins that enter the cell as components of the incoming viral particle and thus occur prior to the formation of phase-separated replication compartments.

Among the known antiviral proteins, a number of “tripartite motif” (TRIM) proteins have been shown to interfere directly with key steps in the life cycles of widely divergent viruses, or exert indirect inhibition as regulators of antiviral signaling (12). While a variety of very distinct mechanisms and functions have been ascribed to TRIM proteins in this context, a characteristic feature of TRIM proteins is their shared architecture. Specifically, TRIM proteins form antiparallel dimers, driven by a central coiled-coil domain, that constitutes one defining feature of the tripartite motif(13–15). At least some TRIM proteins also form higher-order multimers, mediated by interactions between N-terminal RING and/or B-box domains (16, 17), that are also defining features of the tripartite motif. Typically, SPRY or other protein domains situated at the TRIM protein C-terminus enable interactions with viral or cellular targets (12). The propensity of TRIM proteins to form high-order multimeric structures thereby allows polyvalent interactions with targets.

Herein, we describe a loss of function screen to identify ISGs that are mediators of the anti-VSV activity of IFNα. We show that the products of multiple ISGs, including previously unidentified antiviral proteins, contribute to the overall activity of IFNα. Among these proteins, we identify a poorly characterized TRIM protein, TRIM69, as an inhibitor of VSV replication. We show that TRIM69 inhibits VSV replication through a previously unanticipated mechanism of action. Specifically, we find that higher-order TRIM69 multimers target a specific sequence in VSV P. In so doing, TRIM69 inhibits pioneer viral transcription and the formation of VSV replication compartments, resulting in profound reduction of viral RNA synthesis and inhibition of viral replication.

## Results

### Identification of ISGs that mediate the anti-VSV activity of IFNα

To identify genes responsible for the antiviral state, specifically those responsible for inducing VSV (Indiana serotype, VSV_IND_) resistance, we selected a subclone of HT1080 cells in which IFN potently inhibited the replication of a recombinant VSV_IND_ engineered to carry a nanoluciferase (nLuc) reporter gene (VSV_IND_(nLuc)). We also designed an siRNA library containing siRNA pools (Dharamacon SMARTpools containing 4 individual siRNA duplexes) representing the 400 most strongly upregulated genes among a panel of cell lines (SI Apppendix Experimental Procedures), along with 18 siRNA controls (SI Appendix Table S1). We transfected the HT1080 cell line in 96 well plates with the arrayed siRNA library, and treated the cells with 10 U/mL IFNα. The following day, cells were infected at low multiplicity (0.01) with VSV_IND_(nLuc). After a further day, luciferase activity was measured, to identify siRNA pools identified that were able to enhance spreading VSV_IND_ replication in the presence of IFNα (Fig. 1A).

**Fig. 1.**
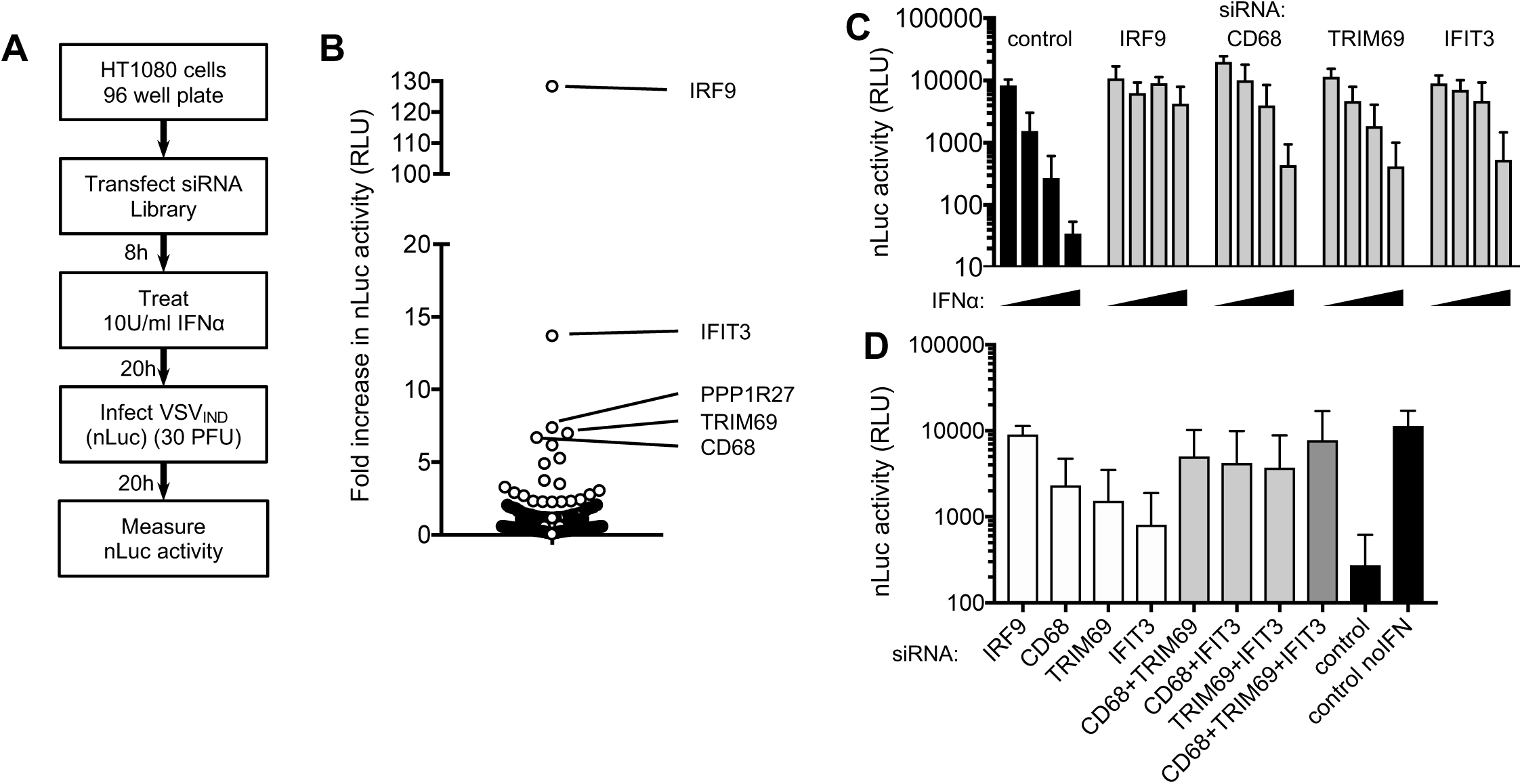
An siRNA based screen for ISGs that inhibit VSV replication. (A) Schematic representation of the screening procedure for siRNA SMARTpools directed against ISGs. (B) VSV_IND_(nLuc) replication (fold increase in nLuc signal compared to controls) in cells transfected with 400 siRNA SMARTpools. (C) Confirmatory assays of VSV_IND_(nLuc) replication (luciferase activity in relative light units (RLU)) in cells transfected with the indicated siRNA SMARTpools and treated with 0, 5, 10 or 20 U/ml of IFNα. (D) VSV_IND_(nLuc) replication (luciferase activity (RLU)) in HT1080 cells transfected with the indicated individual siRNAs, alone or in combination, and treated with 10 U/mL of IFNα.

Using this approach, siRNA pools that increased VSV_IND_(nLuc) replication by >4-fold in both of two separate screens were identified (Fig. 1B). Upon retesting with fresh siRNA pools, IRF9, IFIT3, TRIM69 and CD68 were confirmed as bona fide hits, as the antiviral effect of IFNα was clearly attenuated in siRNA-transfected cells (Fig. 1C). IRF9 was an expected hit, as it is required for type-I IFN signaling, and transfection with siRNAs targeting IRF9 nearly completely abolished the inhibitory activity of IFNα (Fig. 1C). Transfection of each individual siRNAs, or as combinations of two or three siRNAs suggested that IFIT3, CD69 and TRIM69 each contributed to the overall antiviral effect of IFNα (Fig. 1D). Indeed, transfection of all three siRNAs in combination markedly diminished the activity of IFNα (Fig. 1D).

### TRIM69 is an inhibitor of VSV_IND_ replication

The IFIT family of proteins has previously been reported to recognize unusual structures at the ends of viral RNAs (18). We therefore focused our follow-up efforts on CD68 and TRIM69 which had not previously been reported to exhibit antiviral activity. Western blot analysis of HT1080 cells confirmed strong upregulation of both proteins upon IFNα treatment (Fig. 2A). Using a doxycycline-responsive lentiviral expression vector, we established HT1080-derived cell lines that inducibly expressed CD68, TRIM69 or a known anti-VSV protein, Mx1 as a control (SI Appendix Fig. S1A). Doxycycline induction of TRIM69 or Mx1 expression potently inhibited VSV_IND_(nLuc) replication while CD68 expression had only a modest inhibitory effect (Fig. 2B). Profound inhibition of replication was evident when VSV_IND_(nLuc) or unmodified VSV_IND_ was used at low multiplicity to initiate spreading replication assays (Fig. 2B, C). In the latter case, there was a ∼1000-fold reduction in the yield of infectious VSV_IND_ particles following a 26h multi-cycle spreading replication experiment. When VSV_IND_(eGFP) was used in a short-term experiment (MOI=1, 6h, ∼one replication cycle), the number and brightness of GFP+ cells was clearly reduced (Fig. 2D). These data suggest that TRIM69 inhibits a step in the VSV_IND_replication cycle prior to the assembly and release of infectious particles. Consistent with that conclusion, infection of HT1080 cells with VSV_IND_ and western analysis 6h later revealed that VSV_IND_ matrix (M) protein expression was profoundly inhibited by TRIM69 (SI Appendix Fig. S1A).

**Fig. 2.**
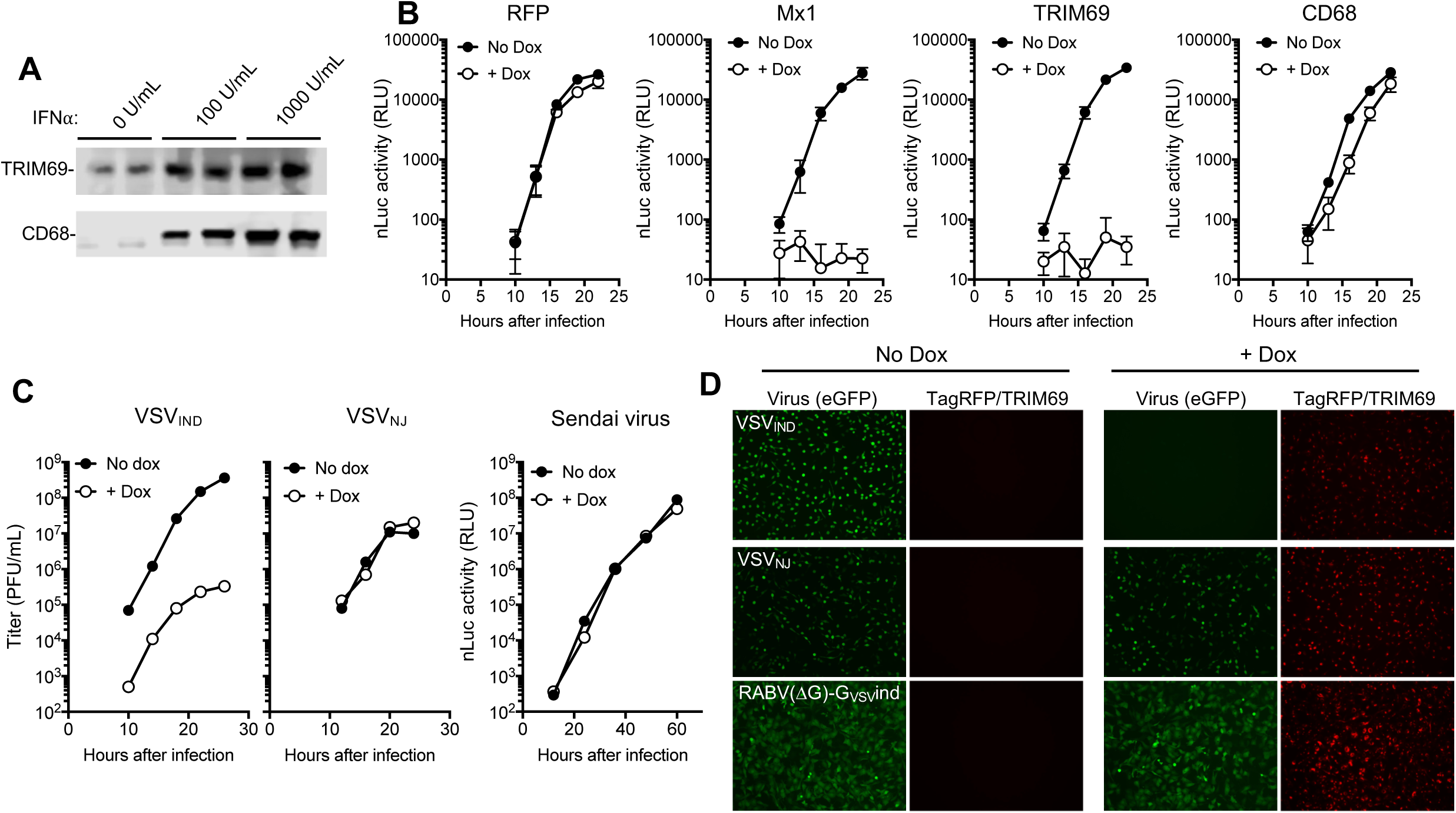
Overexpression of TRIM69 inhibits VSV_IND_ replication. (A) Western blot analysis of endogenous TRIM69 and CD68 expression following IFNα treatment (B) VSV_IND_(nLuc) replication (luciferase activity (RLU)) in HT1080 cells stably transduced with lentiviral vectors containing doxycycline inducible expression cassettes for the indicated genes and infected with 30 PFU VSV_IND_(nLuc) (MOI=0.003). (C) VSV_IND_, VSV_NJ_, and Sendai virus replication in HT1080-TagRFP/TRIM69 cells with or without doxycycline treatment. (D) HT1080-TagRFP/TRIM69 cells treated or not with doxycycline and infected with VSV_IND_(eGFP/P), VSV_NJ_(eGFP) and RabV(eGFP-ΔG)-G_VSVind_ 16h later at MOI=1 for 1 h. Images were acquired at 6 hours post infection (h.p.i).

We tested whether the replication of other distantly or closely related negative-strand RNA viruses were inhibited by TRIM69. Neither Sendai virus, rabies virus or even the New Jersey strain of VSV (VSV_NJ_, that is ∼36% divergent from VSV_IND_) were inhibited by TRIM69 (Fig. 2C, D). Notably, TRIM69 proteins from two divergent mammalian (murine and bovine) species were active inhibitors of VSV_IND_ but TRIM69 proteins from more divergent species; a reptile (chameleon) and fish (Zebrafish) did not inhibit replication (SI Appendix Fig. S1 B,C).

### Anti-VSV_IND_ activity is associated with high-order TRIM69 multimerization

TRIM69 is a member of a large family of TRIM proteins, several of which have been shown to exhibit direct or indirect antiviral activity (12). Like other TRIM proteins, TRIM69 protein forms dimers that is held together by a coiled-coil domain, configured in anti-parallel orientation (15). In addition, for at least some TRIM proteins, the anti-parallel dimers are assembled into higher-order multimeric structures, which are held together via a second dimer or trimer interfaces located in the RING and B-box domains, respectively (16, 17). Among other TRIM proteins, TRIM25 contains a dimeric interface in its RING domain and also shared the highest level of sequence similarity with TRIM69. We used the previously determined crystal structure of a TRIM25 RING domain dimer (17) to identify amino acids at the RING domain dimer interface (V95, L96 and L99 in TRIM69, analogous to V68, L69 and V72 in TRIM25, Fig. 3A, SI Appendix Fig.S2A) whose mutation might disrupt the TRIM69 RING dimer interface, and, perhaps, higher-order TRIM69 multimerization (Fig. 3A).

**Fig. 3.**
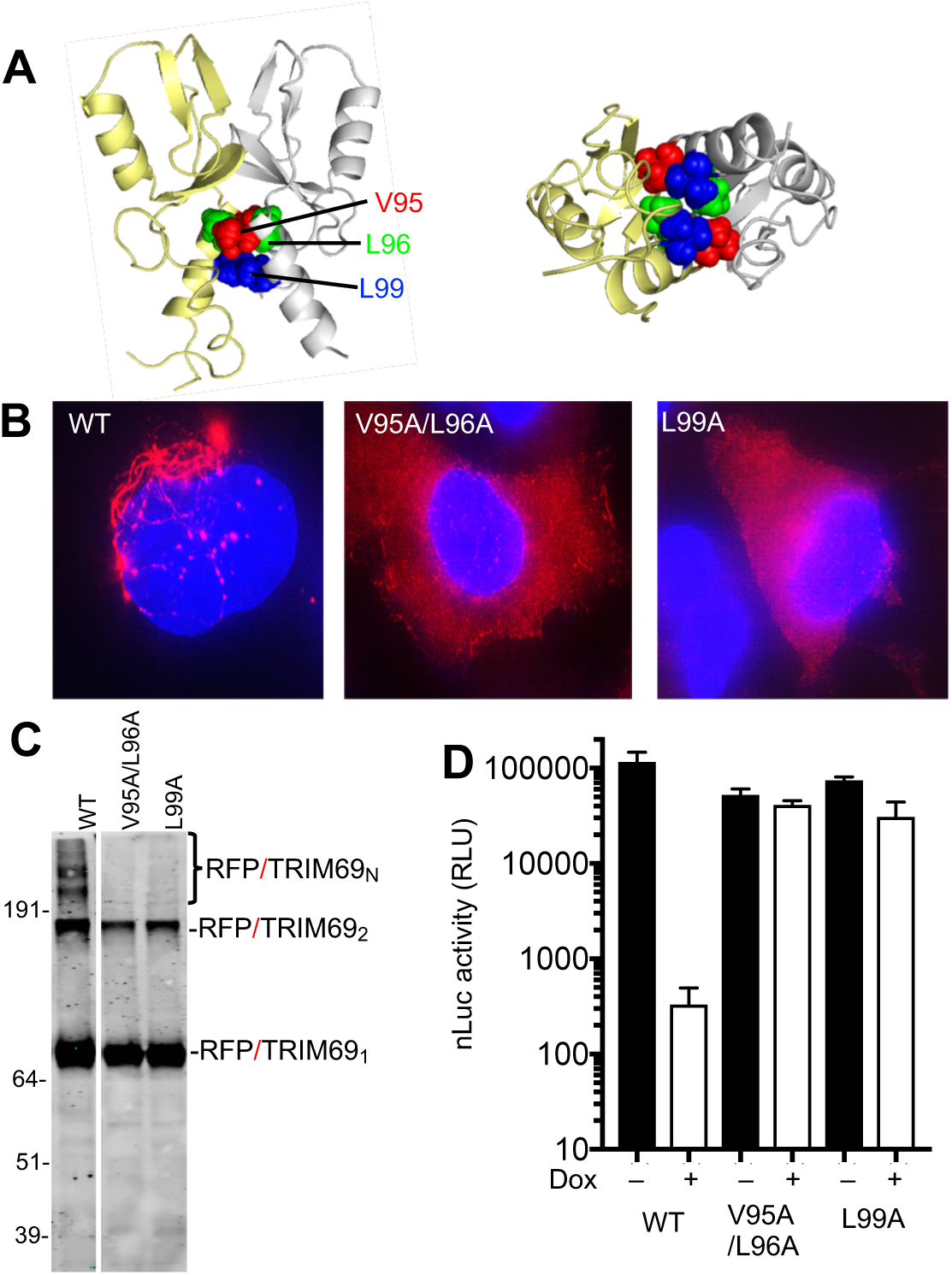
Effects of TRIM69 multimerization on antiviral activity. (A) Crystal structure of dimeric TRIM25 RING domain indicating amino acids at the dimer interface: V68 (red), L69 (green) and V72 (blue) analogous to V95, L96 and L99 in TRIM69. (B) Deconvolution microscopic images of WT and mutant TagRFP/TRIM69 fusion proteins expressed in doxycycline inducible HT1080 cells (C) Western blot analysis of WT and mutant TagRFP/TRIM69 fusion proteins following treatment of cells with EGS crosslinker prior to cell lysis (D) VSV_IND_(nLuc) replication (luciferase activity (RLU)) in HT1080 cells stably transduced with lentiviral vectors contain doxycycline inducible expression cassettes for WT and mutant TagRFP/TRIM69 proteins.

Deconvolution microscopic imaging of TagRFP/TRIM69 expressed in HT1080 cells (Fig. 3B) as well as 3D structured illumination (3D-SIM) super-resolution imaging of mScarlet/TRIM69 (SI Appendix Fig. S2B), showed that unmodified TRIM69 formed filamentous structures and cytoplasmic bodies in the cytoplasm of cells, consistent with the finding that it assembles into high-order structures. Moreover, western analysis of cell lysates generated after treatment of cells with a protein crosslinker (EGS) showed that TRIM69 formed dimers and, apparently, higher-order multimers (Fig. 3C). Notably, mutation of the predicted RING domain dimer interface (V95A/L96A double mutant or L99A single mutant) resulted in TagRFP/TRIM69 proteins that formed crosslinkable dimers, but not higher order multimers (Fig. 3C), and exhibited a diffuse rather than filamentous distribution in the cell cytoplasm (Fig. 3B). Notably, V95A/L96A and L99A mutations abolished the ability of TRIM69 to inhibit VSV_IND_(nLuc) replication, suggesting that higher-order multimerization is necessary for antiviral activity (Fig. 3D).

### Recruitment of TRIM69 to sites of VSV replication and inhibition of replication compartment formation

Like many RNA viruses, VSV partitions its replication machinery into specialized compartments within which RNA synthesis occurs (6–9). In the case of VSV, recent work has shown that these compartments are liquid inclusions that are not bounded by membranes, but instead exhibit characteristics of phase-separation (10). Replication compartments are conveniently labelled and localized using VSV clones modified to append a fluorescent protein to the amino terminus of P. We infected cells with VSV_IND_(NeonGreen/P) and visualized replication compartments using NeonGreen, along with fluorescence in situ hybridization (FISH) probes directed to the negative strand viral RNA. This analysis showed that the presence of TRIM69 profoundly attenuated the formation of replication compartments marked by the P protein and negative strand RNA (SI Appendix Fig. S3). Additionally, 3D-SIM imaging of cells expressing mScarlet/TRIM69 and infected with VSV_IND_(NeonGreen/P) revealed that the smaller P accumulations that were observed in TRIM69 expressing cells were typically colocalized with the mScarlet/TRIM69 filaments. Indeed, many of the P accumulations appeared to adopt an elongated, almost filamentous structure, different in shape and size to the compartments typically observed in VSV infected cells, and coincident with TRIM69 filaments (SI Appendix Fig. S4). Overall, these data suggested that a viral or cellular component governing formation of viral replication compartments associates with TRIM69 and that this association ultimately inhibits replication compartment formation.

### The VSV phosphoprotein (P) is the viral determinant of TRIM69 sensitivity

To determine how TRIM69 inhibits VSV replication, we selected mutant VSV derivatives that were resistant to TagRFP/TRIM69. We infected on HT1080 cells expressing TagRFP/TRIM69 infected at high multiplicity (MOI =10) with VSV_IND_(eGFP), or a VSV_IND_ clone which eGFP was appended to P (VSV_IND_(eGFP/P)). Three growing plaques each were picked for VSV_IND_(eGFP) and VSV_IND_(eGFP/P), and amplified on HT1080 cells expressing TagRFP/TRIM69. These viruses infected HT1080 cells with equivalent efficiency, whether or not TagRFP/TRIM69 was induced (Fig. 4A, SI Appendix Fig S5A). Sequencing of the viral genome of these TRIM69-resistant viruses (TR), showed the concentration of nonsynonymous mutations within a short peptide sequence (amino acids 66-71) within P (Fig. 4B, SI Appendix Fig S5B). For 4/6 viruses, the only nonsynonymous mutation found in the genome is located within this peptide thus indicating these mutations are responsible for the resistance to TRIM69. This peptide sequence is within a region of P that contacts the globular connector, methyl transferase and CTD of L, and is also responsible for stimulating polymerase activity (the L-stimulatory region, or LSR, Fig. 4B) (19, 20). Although located in a crucial region for efficient viral RNA synthesis, the TR mutations did not affect replication of VSV_IND_(eGFP) or VSV_IND_(eGFP/P) in Vero cells (SI Appendix Fig S5C).

**Fig. 4.**
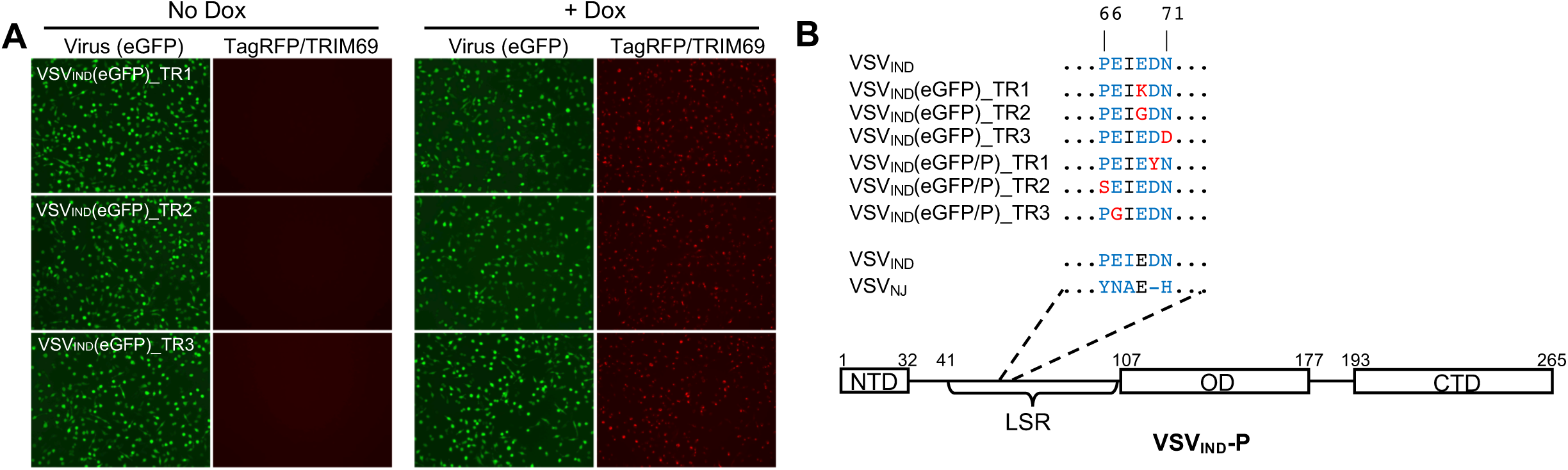
The VSV-P protein determines sensitivity to TRIM69. (A) HT1080-TagRFP/TRIM69 cells treated or not with doxycycline and infected with VSV_IND_(eGFP/P) (TR1-3) 16h later at MOI=1 for 1 h. Images were acquired at 6 (h.p.i) (B) Schematic of VSV_IND_ P showing its modular organization into three domains; N-terminal domain (NTD), oligomerization domain (OD) and C-terminal domain (CTD). The L stimulatory region (LSR, aa 41-106) and the peptide containing the point mutations found in TRIM69-resistant clones (aa 66-71) is indicated. Point mutations are shown in red and residues mutated at least once in TR viruses are colored in blue.

### TRIM69 associates with VSV_IND_ P

We next investigated how the TR mutations in P exerted their effects. While WT TagRFP/TRIM69 effectively prevented the formation of VSV_IND_(eGFP/P) replication compartments, the two VSV_IND_(eGFP/P) TR mutants (D70Y and E67G) formed prominent replication compartments that showed no association with TRIM69 filaments or accumulations (SI Appendix Fig. S6). Strikingly, the multimerization-defective mutants of TRIM69 (V95A/L96A and L99A) that did not exhibit antiviral activity and were ordinarily diffusely distributed in the cytosol (Fig 3B), were nearly completely relocalized to replication compartments in VSV_IND_(NeonGreen/P) or VSV_IND_(eGFP/P) infected cells (Fig. 5A, SI Appendix Fig. S7). Thus, TRIM69 recruitment to replication compartments was not itself sufficient to inhibit VSV_IND_ replication. Moreover, the dramatic redistribution of TRIM69 (L99A) to replication compartments in VSV_IND_ infected cells was completely absent in VSV_IND_(eGFP/P)_TR1(D70Y) and VSV_IND_(eGFP/P)_TR3(E67G) infected cells. Together, these data indicate that specific sequences with P are necessary for the recruitment of TRIM69 to the VSV_IND_ replication machinery and vice versa. To determine whether P was sufficient for TRIM69 recruitment, we expressed eGFP/P by transfection, in the absence of any other viral proteins. In contrast to the situation in VSV_IND_ infected cells, eGFP/P expressed alone was diffusely distributed throughout the cytoplasm. However, in TagRFP/TRIM69 expressing cells, eGFP/P was recruited to the TagRFP/TRIM69 accumulations and the two proteins colocalized extensively (SI Appendix Fig. S8A, B). Again, this colocalization was dependent on the viral determinant of TRIM69 sensitivity, as there was no colocalization between TagRFP/TRIM69 and eGFP/P(E67G).

**Fig. 5.**
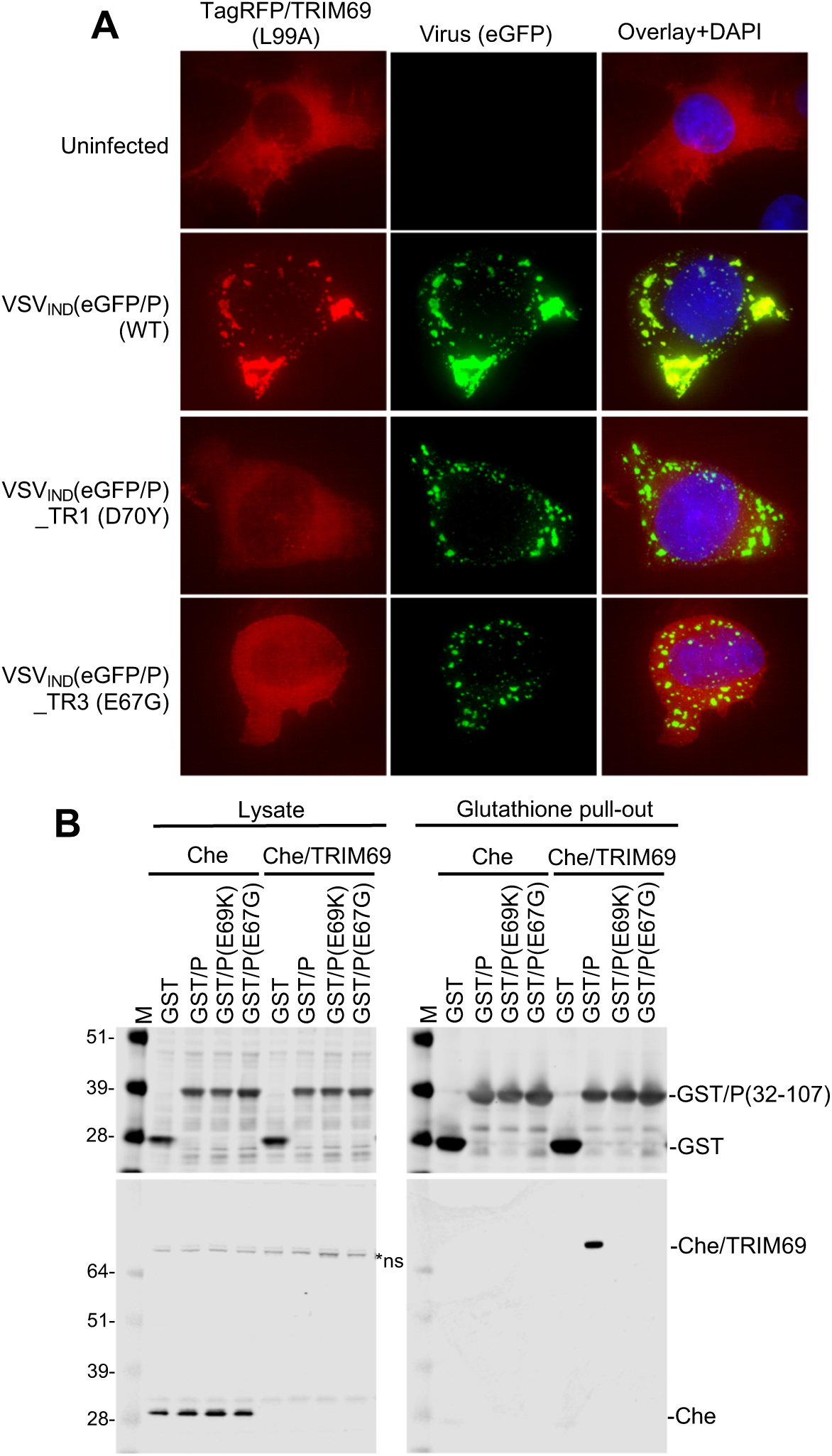
TRIM69 binds to P. (A) Deconvolution microscopic images of HT1080-TagRFP(L99A) mutant cells infected with WT or mutant VSV_IND_(eGFP/P). (B) Western blot (anti-GST, upper panels and anti-Cherry fluorescent protein, lower panels) analyses of cell lysates and glutathione pull-out fractions from 293T cells transfected with plasmids expressing Cherry/TRIM69 and either WT or mutant GST-P proteins.

To determine whether P physically associated with TRIM69, we generated glutathione-S-transferase (GST) proteins fused to a region of P encompassing the LSR domain (amino acids 32-107). We also constructed GST/P fusion protein containing mutant LSR domains from VSV_IND_(eGFP)_TR1 (E69K) and from VSV_IND_(eGFP/P)_TR3 (E67G) (Fig.4B). The GST-P fusion proteins were coexpressed with Cherry/TRIM69 in 293T cells and then purified from cell lysates with glutathione Sepharose. Cherry/TRIM69 was nearly undetectable in clarified cell lysates, presumably due to its propensity to form higher-order multimers that were poorly soluble in non-denaturing detergents (Fig. 5B). Nevertheless, GST/P coprecipitated Cherry/TRIM69, such that it was highly enriched in precipitated fractions (Fig. 5B). Conversely, GST/P proteins bearing the TR mutations (E69K or E67G) did not precipitate detectable amounts of Cherry/TRIM69 (Fig. 5B). Thus, the isolated LSR domain was sufficient for the physical association of P with TRIM69.

### Mechanism of VSV replication inhibition by TRIM69

The above data provides evidence that TRIM69 physically associates with P and ultimately inhibits VSV_IND_ replication compartment formation. However, the assembly of replication compartments requires multiple prior steps, each of which are potential targets of TRIM69. For example, because TRIM proteins can act as ubiquitin ligases(12), it was conceivable that TRIM69 might mediate destruction of one or more viral proteins. Alternatively, interaction with P could block replication through a non-degrative mechanism by inhibiting: (1) an initial ‘pioneer’ round of transcription that employs the incoming virion RNA genome as a template to generate viral mRNAs, (2) the translation of these new viral mRNAs to generate N, P and L proteins and the assembly of newly synthesized N, P and L proteins with full-length negative strand RNAs.

To determine whether TRIM69 targeted incoming viral proteins for degradation, we first infected cells at high MOI with ^35^S labelled virions and monitored the levels of virion proteins in the presence of cycloheximide (CHX) to prevent new protein synthesis and VSV replication. No substantial difference in the decay of incoming viral proteins in the presence or absence of TRIM69 was detected (SI Appendix Fig S9A). Next, we monitored the levels of P protein following transfection into cells in which TRIM69 expression was, or was not, induced. Despite obvious recruitment of EGFP/P to sites of TRIM69 concentration (SI Appendix Fig S8), overall levels of WT and TR mutant EGFP/P were equivalent and unaffected by TRIM69 (SI Appendix Fig S9B). Finally, we found that WT and as well as active (K82A) and inactive (L99A) mutants of TRIM69 exhibited approximately equivalent levels of autoubiquitination (SI Appendix Fig S9C). Overall, we found no evidence the TRIM69 drives VSV protein degradation, and the ubiquitin ligase activity of TRIM69 was unable to account for its antiviral activity.

Next, we quantified new viral mRNA (pioneer) transcription from incoming virion RNA templates in cells treated with CHX to prevent protein synthesis and replication. First a single molecule FISH assay and a pool of oligonucleotide probes directed at the plus strand N mRNA demonstrated a clear reduction in pioneer N mRNA transcription in cells expressing TRIM69 (Fig. 6A, B). In an alternative approach to measure VSV transcription, we labelled target cells with ^32^P orthophosphate, in the presence of actinomycin D (to block host mRNA synthesis). At 5h after VSV_IND_(eGFP/P) infection, transcripts corresponding to L, G, and M mRNAs were clearly detectable (Fig. 6C). Transcripts corresponding to N and/or eGFP/P mRNAs were also detected but could not be distinguished from each other due to co-migration. As expected, in the absence of CHX, mRNA synthesis and genome replication of VSV_IND_(eGFP/P) were inhibited by TRIM69 but not TRIM69(L99A), while VSV_IND_(eGFP/P)_TR3 mRNA synthesis and genome replication was insensitive to TRIM69. Importantly, when CHX was used to prevent protein synthesis and RNA replication, the presence of TRIM69 reduced the levels of all nascent VSV_IND_ mRNA transcripts (Fig. 6C). The magnitude of the effect on transcript levels appeared greatest for the L mRNA, least for the N and P mRNAs and of intermediate magnitude for M and G mRNAs (Fig. 6C, SI Appendix Fig. S9B). The multimerization-defective TRIM69 (L99A) mutant did not affect pioneer mRNA synthesis (Fig. 6C). Thus, TRIM69 inhibited VSV_IND_transcription, with apparently greater effect on genes encoded near the 5’ end of the negative strand genome.

**Fig. 6.**
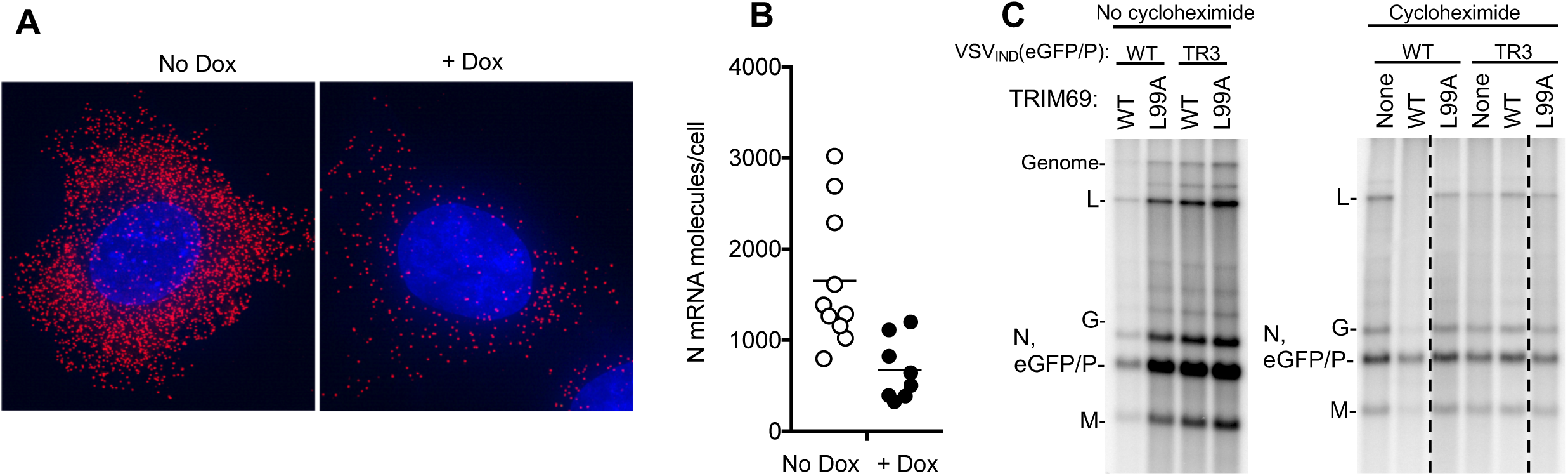
TRIM69 inhibits VSV_IND_ transcription. (A) Single molecule FISH analysis of HT1080-TRIM69/myc cells treated with cycloheximide and infected with VSV_IND_(GFP/P) for 3h, using probes directed to N mRNA. Individual representative cells are shown. (B) Single molecule FISH analysis of HT1080-TRIM69 as in A, each symbol represents an individual cell and the number of mRNA molecules in each cell is plotted. Unpaired *t* test, p=0.0036 (C) HT1080 cells expressing WT or mutant (L99A) TagRFP/TRIM69 and cultured with actinomycin-D and ^32^P orthophosphate, were infected with VSV_IND_(eGFP/P) or VSV_IND_(eGFP/P) (E67G) at MOI=100. To prevent genome replication, cells were also treated with cycloheximide as indicated. RNA was extracted at 5 h.p.i. and analyzed on an agarose-urea gel. Note that N and eGFP/P mRNAs comigrate.

## Discussion

A number of ISGs including Mx1, PKR, IFITM3, and tetherin have previously been reported to inhibit VSV replication (21–24). However, it was not known whether this list represents a complete catalogue of ISGs proteins with anti-VSV activity. We found that a number of antiviral ISGs contribute to the induced antiviral state that prevents VSV_IND_ replication in IFNα treated cells. Among these, we found that TRIM69 has a previously undescribed mechanism of action, inhibiting VSV_IND_ transcription by targeting the polymerase cofactor, P.

TRIM69 joins a growing list of TRIM proteins that have been shown to exhibit antiviral activity through various mechanisms (12). During the course of this work, TRIM69 itself was reported to inhibit Dengue virus replication, albeit via a different mechanism to that described herein, namely ubiquitin-induced degradation of the viral NS3 RNA helicase (25). Other examples of antiviral TRIM proteins include TRIM5*α*, which inhibits early stages of retroviral infection by binding in a polyvalent manner to incoming retroviral capsids, promoting premature uncoating and degradation of virion components (26, 27). Another TRIM protein, TRIM25 has been reported to promote ubiquitination of the RNA sensor RIG-I thereby inducing binding to mitochondrial antiviral signaling (MAVS) protein and stimulation of IFN production (28), although this model has recently been challenged (29). TRIM25 is also an important cofactor of the zinc finger antiviral protein (ZAP), that senses and depletes CG-rich viral RNAs, although the mechanism by which enables ZAP activity remain unclear (30). A variety of other TRIM proteins have been reported to inhibit viral replication directly or indirectly through less well characterized mechanisms (12).

In the two aforementioned examples, high order multimerization is crucial for activity. TRIM5 higher-order multimerization, driven by a B-box domain facilitates the formation of a hexagonal lattice on the surface of incoming retroviral capsid, enabling polyvalent interaction between the capsid hexagonal lattice and a complementary hexagonal TRIM5 lattice (16, 31). In this case, higher-order multimerization results in a more avid interaction between TRIM5 and is viral capsid protein target. For TRIM25, RING domain dimerization enables engagement of ubiquitin conjugated E2 enzymes and higher-order assembly of the RIG-I signalosome (17). Herein, we found that the RING domain dimer interface, analogous to that found in TRIM25 was required for higher-order TRIM69 multimerization, the formation of TRIM69 filaments and antiviral function. However, abolition of high-order TRIM69 multimerization by mutation of the RING domain dimer interface did not prevent recruitment into VSV_IND_ replication compartments. Rather, recruitment of dimeric TRIM69 to replication compartments remained efficient, but was inconsequential to VSV replication. Thus, for TRIM69, RING domain-mediated multimerization appeared to be required for antiviral activity, but not target recognition. As RING domain dimerization might lead to E2 recruitment as well as high order multimer formation, it is not clear whether higher order multimerization *per se*, or downstream E2 recruitment is essential for TRIM69 activity. However, the lack of effect of TRIM69 on incoming virion protein stability, or on co-expressed P levels, coupled with the finding that the L99A mutant maintained ubiquitin ligase activity argues that destruction of virion proteins is not central to the mechanism of action of TRIM69. Unfortunately, we were not able to identify a TRIM69 mutant that maintained higher-order multimer formation, but abolished ubiquitination activity.

We did not formally demonstrate that TRIM69 directly binds to P and it is possible that P interacts with some bridging host protein(s) that is(are) bound by TRIM69. However, a direct interaction between TRIM69 and the LSR domain of P is the most likely molecular event underlying recognition and disruption of the viral transcription/replication machinery. P is required for the interaction between L and the N-coated RNA template (19, 32), and thus for initial transcription following viral entry, as well as for the formation of replication compartments (10). Given that P plays a pivotal, multifactorial role in VSV RNA synthesis and has no cellular homologs it represents an attractive target for intrinsic immune defenses. Moreover, the pioneer round of transcription may represent a point of vulnerability where the number of virion targets is low, at which ISG mediated inhibition might exert maximal effects. Nevertheless, with the caveat that overexpressed TRIM69 was used, we noted the formation of elongated filamentous accumulations of P, coincident with TRIM69 filaments in TRIM69 expressing cells, rather than the spherical droplets that normally characterize the phase-separated VSV replication compartments. This suggests that TRIM69 might inhibit replication compartment formation in addition to its effects on pioneer transcription. Because the L99A TRIM69 mutant retained the ability to be recruited by P and localizes to the phase-separated replication compartments yet did not inhibit VSV_IND_ transcription, it appears that the interaction of TRIM69 with the LSR of P does not prevent functional P-L complex formation. While further study will be required to elucidate the molecular details of how TRIM69 recognizes and disrupts the VSV replication machinery, these findings reveal a new facet of the diverse ways in which in which IFNs control the replication of viruses.

## Acknowledgements

We thank members of the Bieniasz and Hatziioannou labs for helpful discussions and advice and the MicRoN (Microscope Resources on the North Quad) core for their assistance. We thank Rachel Liberatore and Benhur Lee for reagents. This work was supported by grants from the NIH R37AI64003 (to P.D.B.), R37AI059371 (to S.P.J.W.), and R01GM112508/AI150479 (to O.P.)

## SI Appendix

### SI Figure legends

**Fig S1.**
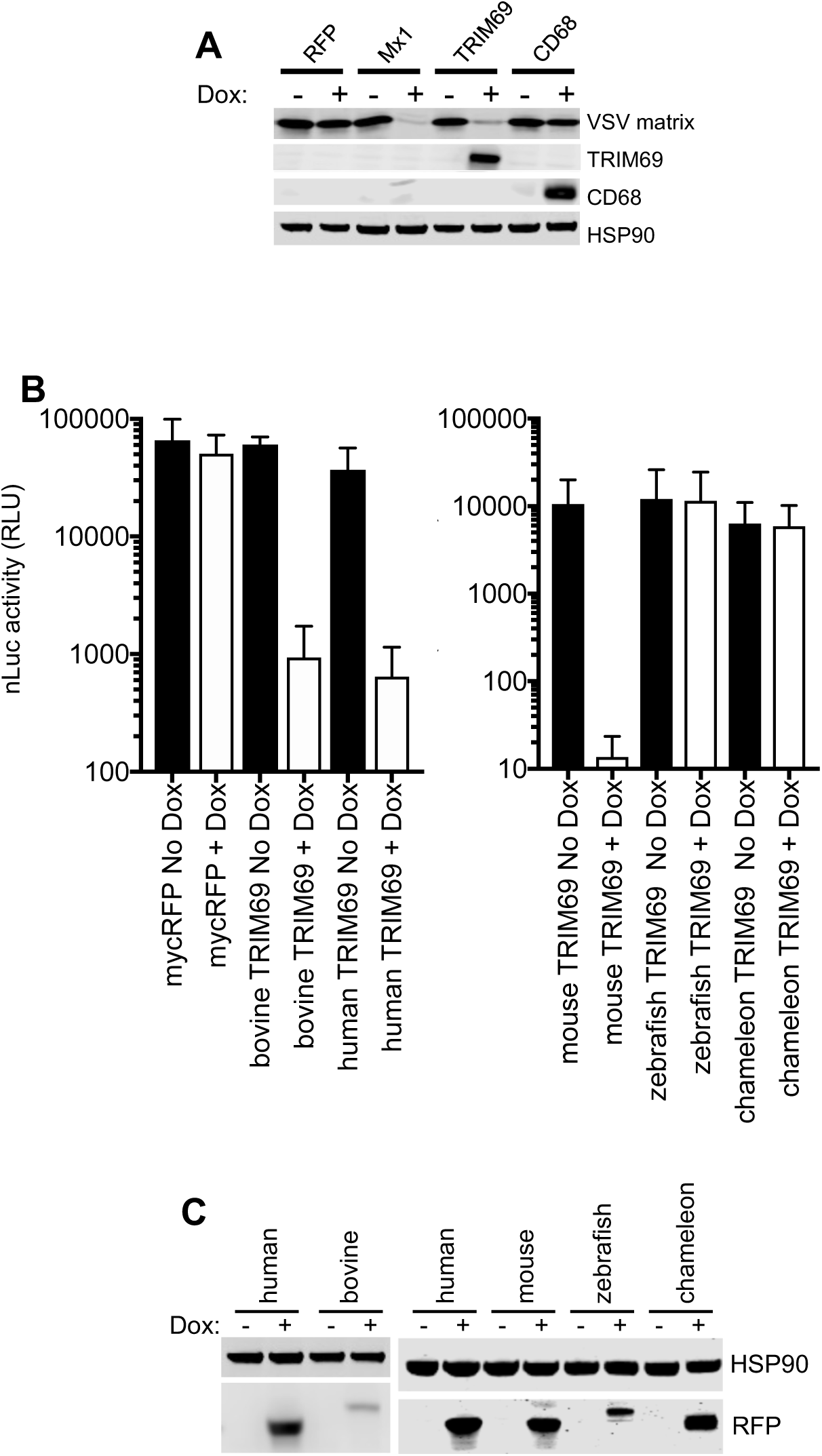
(A) Western blot analysis of VSV_IND_ M, TRIM69, CD68 expression following induction with Dox and infection with VSV_IND_ for 5h (B) VSV_IND_(nLuc) replication in HT1080 cells expressing doxycycline inducible myc-tagged TRIM69 proteins form various species (C) Western blot analysis of TRIM69-myc protein expression following induction with doxycycline

**Fig. S2.**
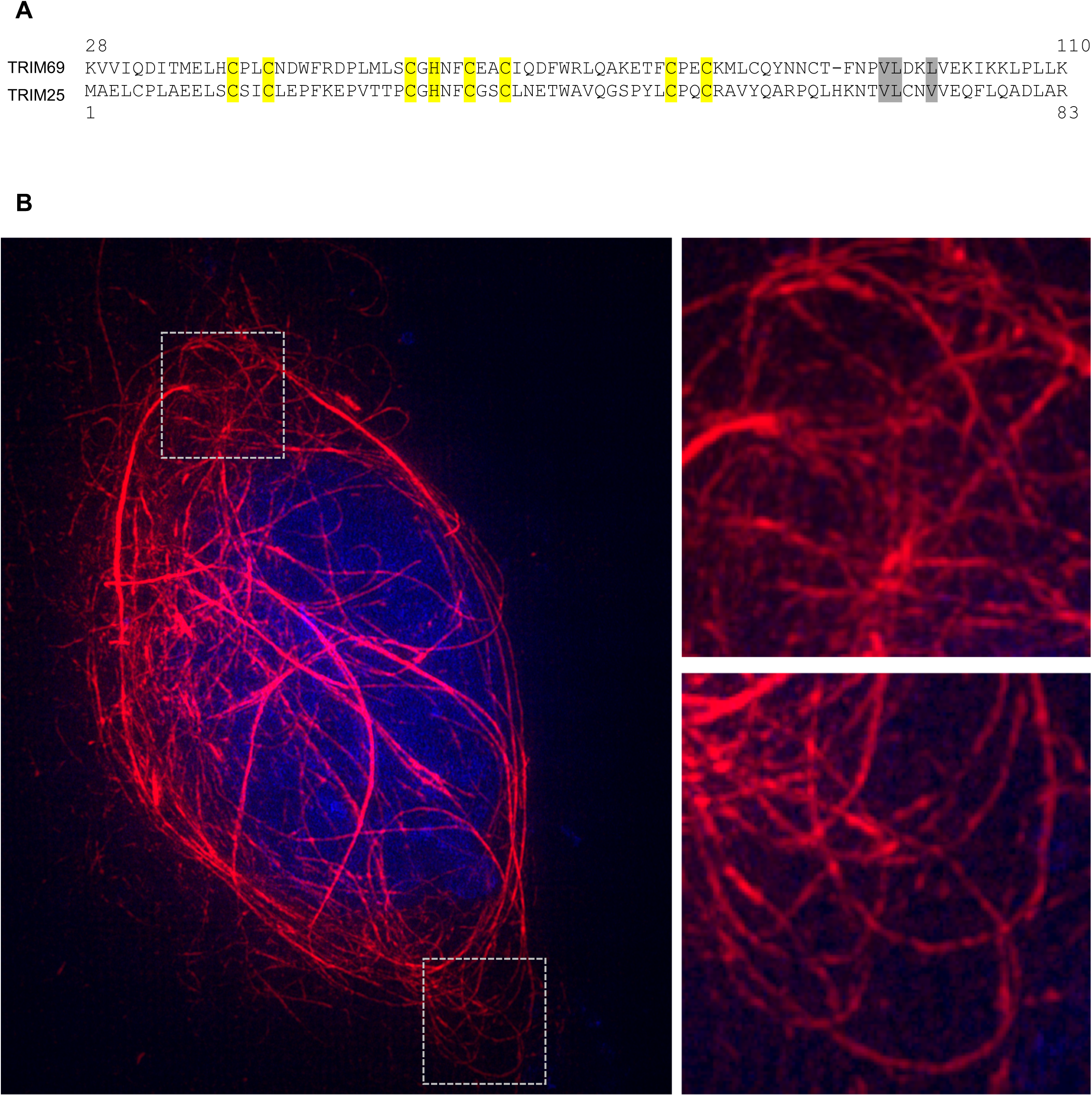
(A) Alignment of TRIM69 and TRIM25 RING domain. Yellow= Zinc coordinating residues. Grey = mutated residues at the dimer interface. (B) 3D SIM images of a mScarlet-TRIM69 expressing cell, with expanded views of the boxed areas.

**Fig. S3.**
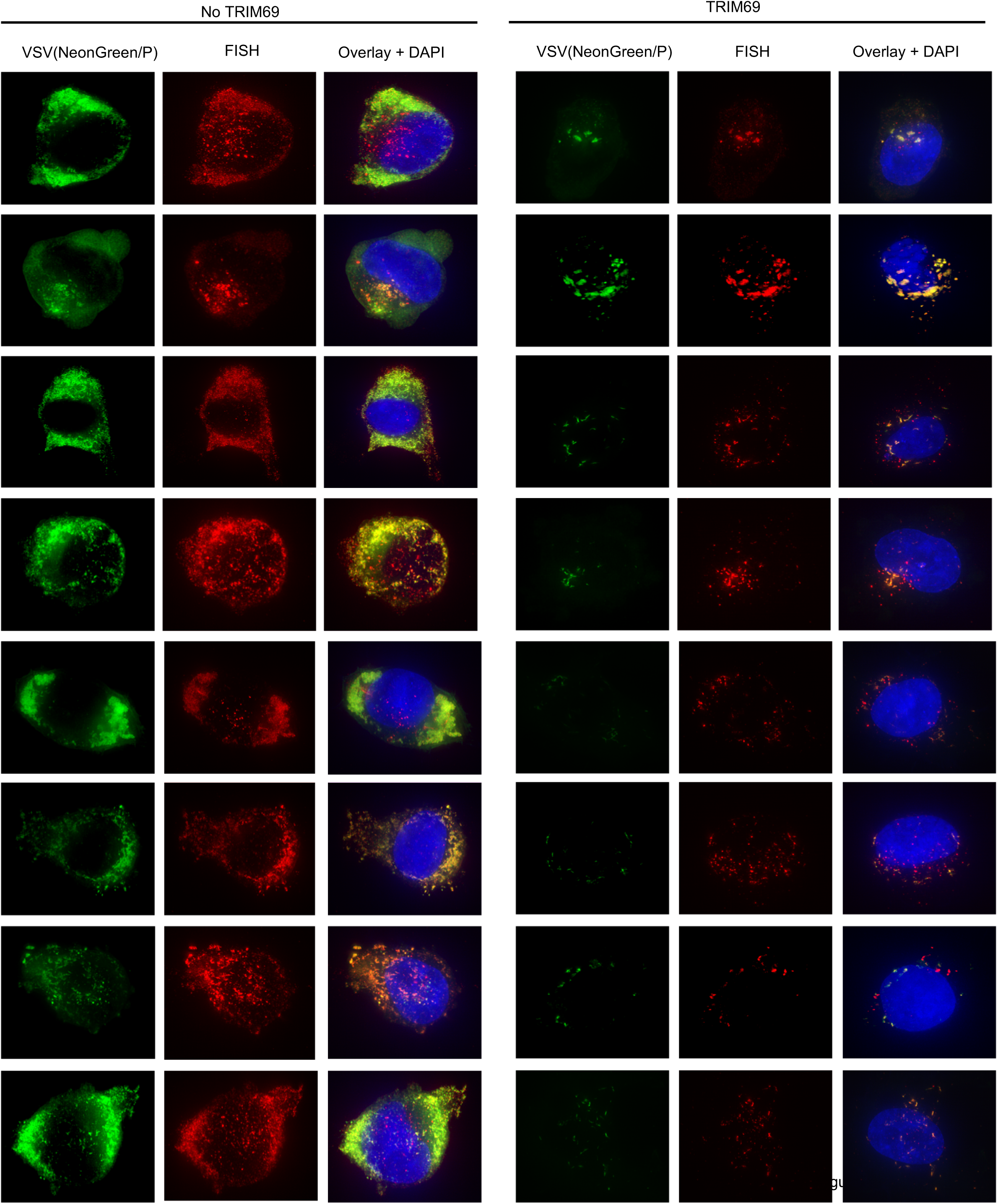
Gallery of randomly selected cells expressing (right) or not expressing (left) TRIM69, fixed 4h after infection with VSV_IND_(NeonGreen/P) and subjected to FISH with probes targeting the negative strand RNA (N gene)

**Fig. S4.**
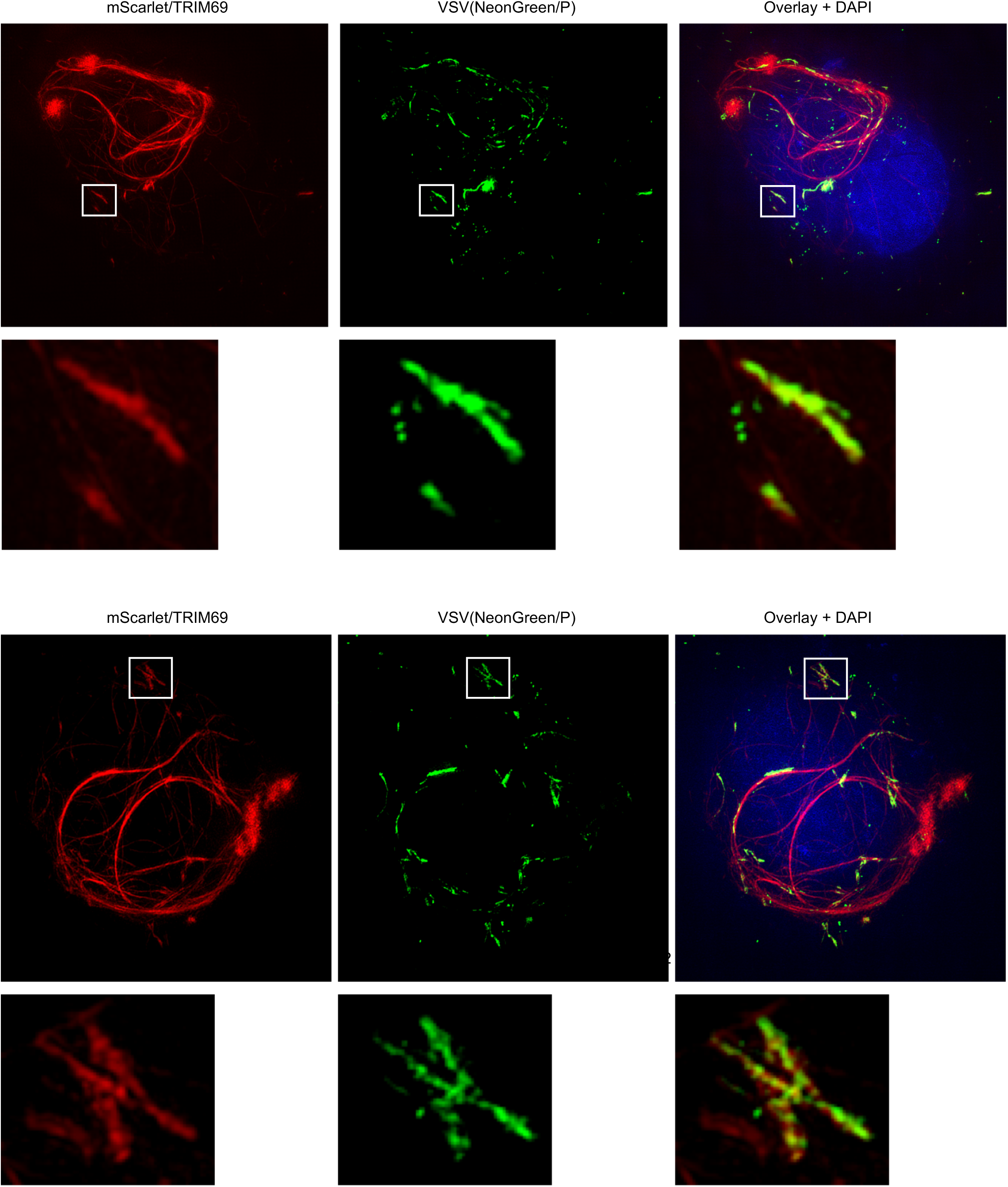
3D SIM images of two mScarlet-TRIM69 expressing cells infected with VSV_IND_(NeonGreen/P), with expanded views of the boxed areas.

**Fig. S5.**
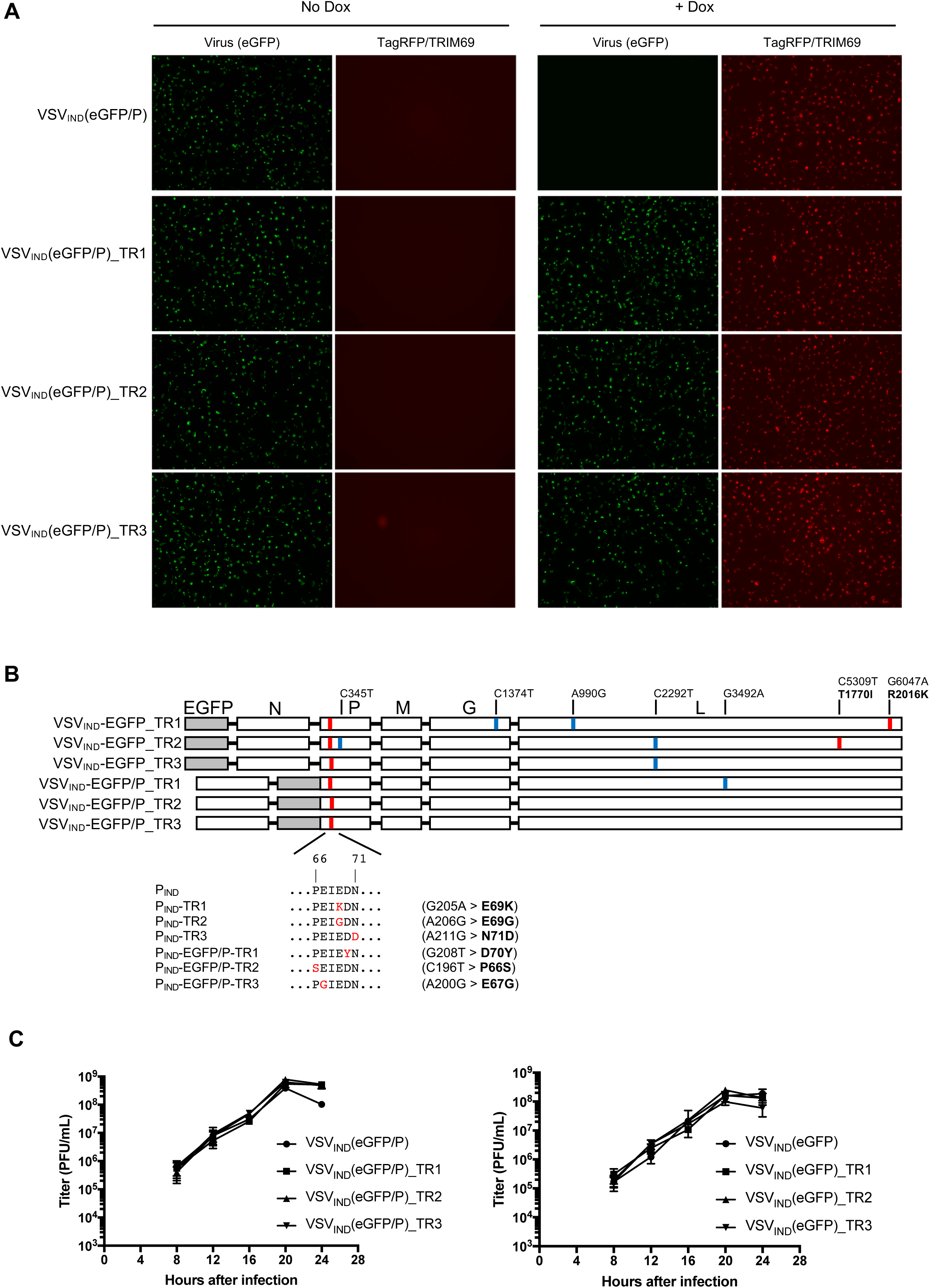
(A) HT1080-TagRFP/TRIM69 cells were seeded and simultaneously treated or not with doxycycline. Sixteen hours later, cells were infected with VSV_IND_(eGFP/P) and TRIM69-resistant (TR) clones at MOI=1 for 1 h. Images were acquired at 6 h.p.i. (B) Complete genome sequences of TRIM69 resistant viruses. Synonymous substitutions are indicated in blue, non-synonymous mutations are indicated in red. All TR viruses contained a non-synonymous mutation in the P (aa 66-71) peptide (C) Replication of WT and TR viruses in Vero cells, infected at an MOI of 0.05

**Fig. S6.**
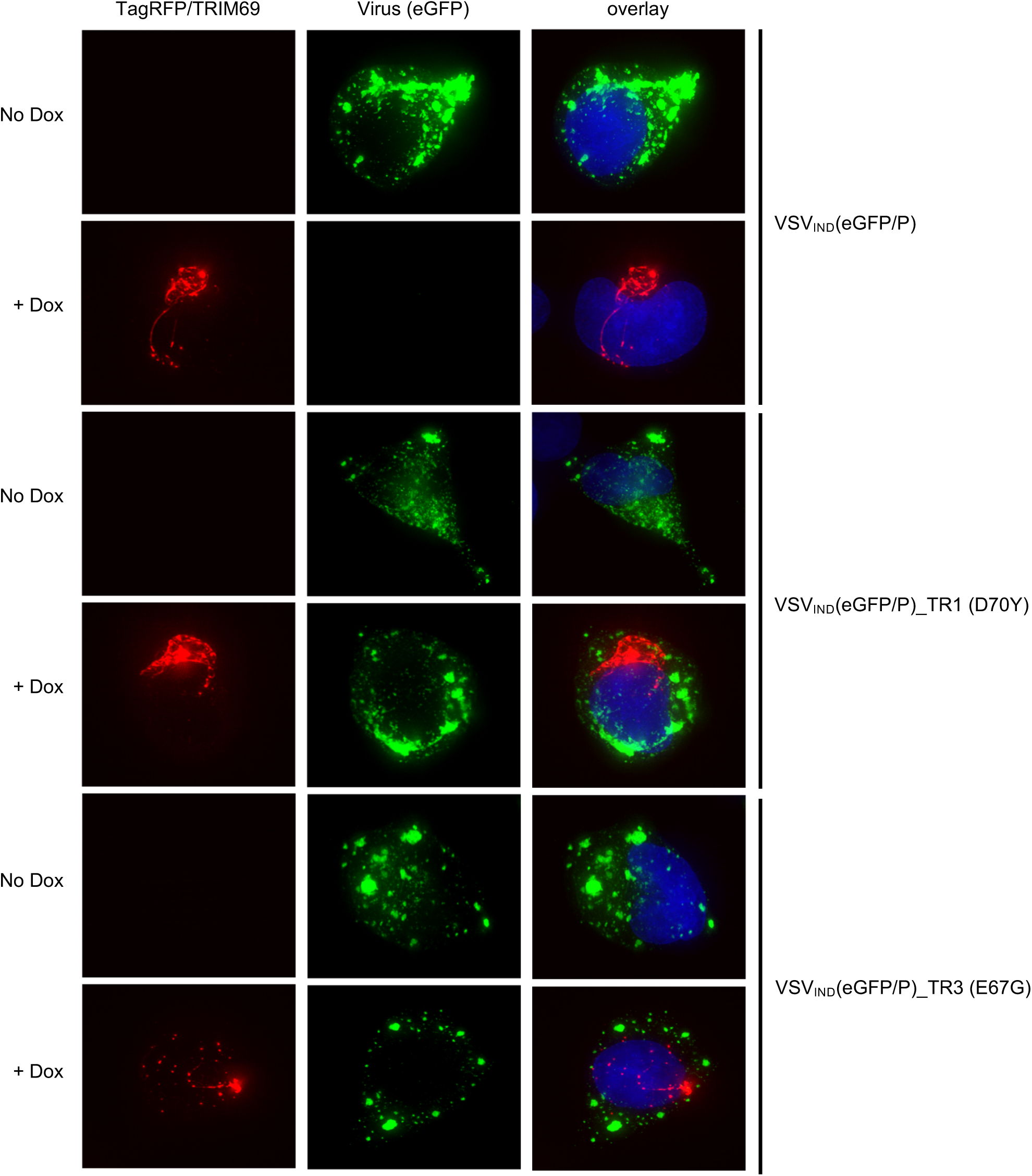
HT1080-TagRFP/TRIM69 cells were treated or not treated with doxycycline and infected with VSV_IND_(eGFP/P) and TRIM69-resistant mutants (D70Y) or (E67G). Images were acquired at 4 h.p.i.

**Fig. S7.**
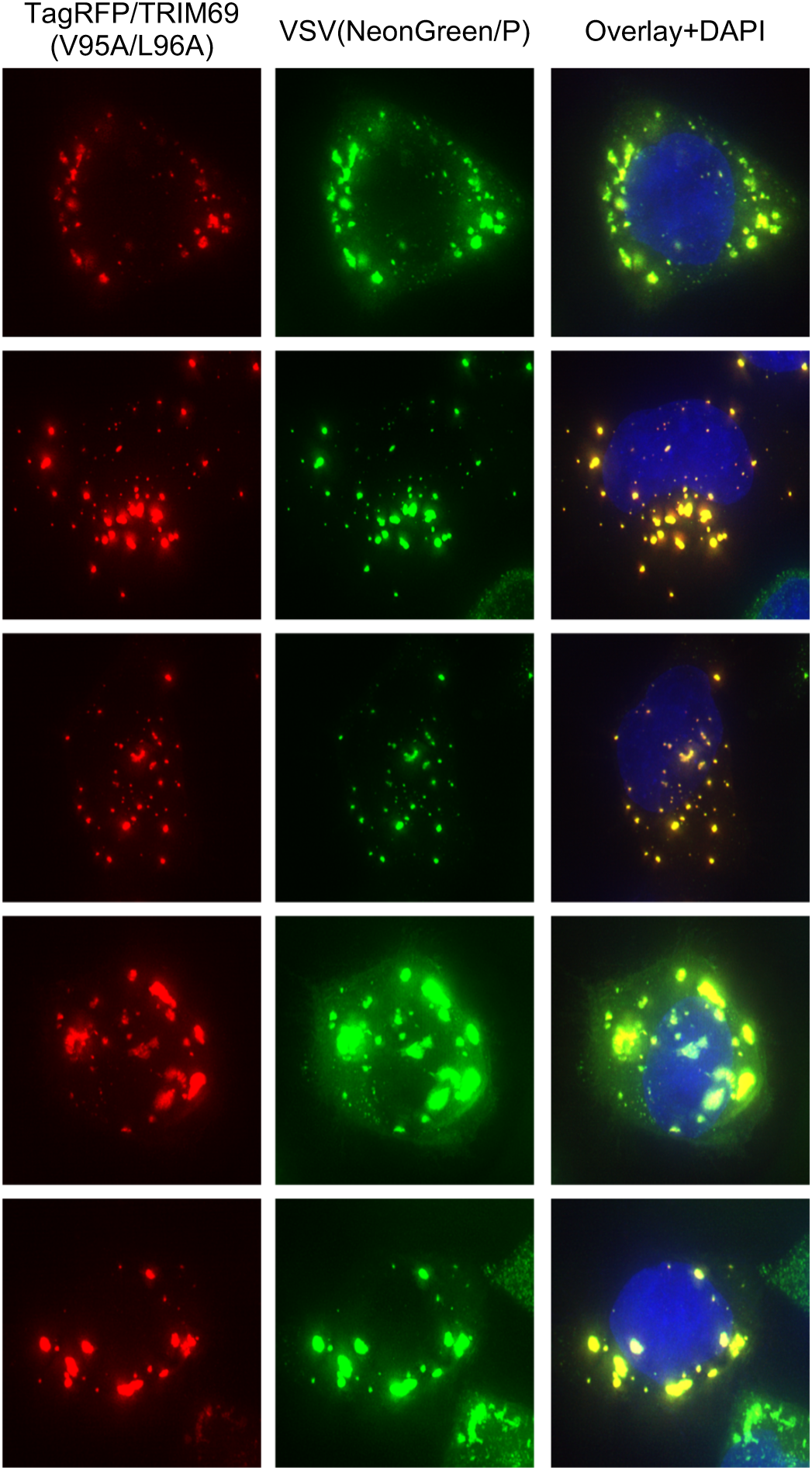
Gallery of HT1080-TagRFP/TRIM69 (V95A L96A) cells treated with doxycycline and infected with VSV_IND_(NeonGreen/P). Images were acquired at 4 h.p.i.

**Fig. S8.**
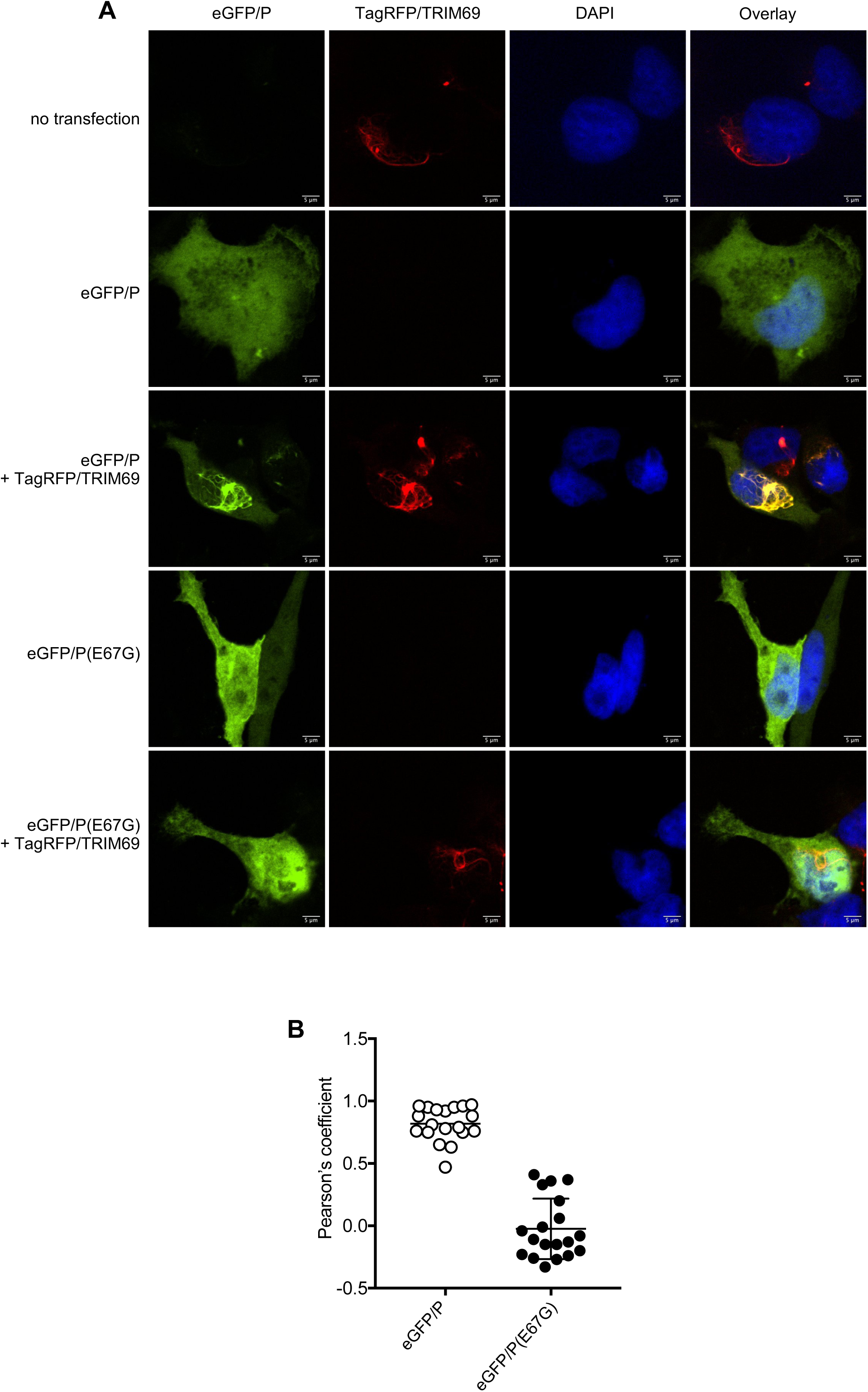
(A) HT1080-TagRFP/TRIM69 cells were seeded and simultaneously treated or not with doxycycline. Sixteen hours later, cells were transfected or not with plasmids expressing eGFP/P or eGFP/P(E67G). Twenty-four hours post-transfection, cells were fixed, stained with DAPI and imaged with a spinning disk confocal microscope. Scale bar, 1 μM. (B) Colocalization analysis of eGFP/P and TagRFP/TRIM69 using ImageJ software. Unpaired *t* test, n=19, P<0.0001.

**Fig. S9.**
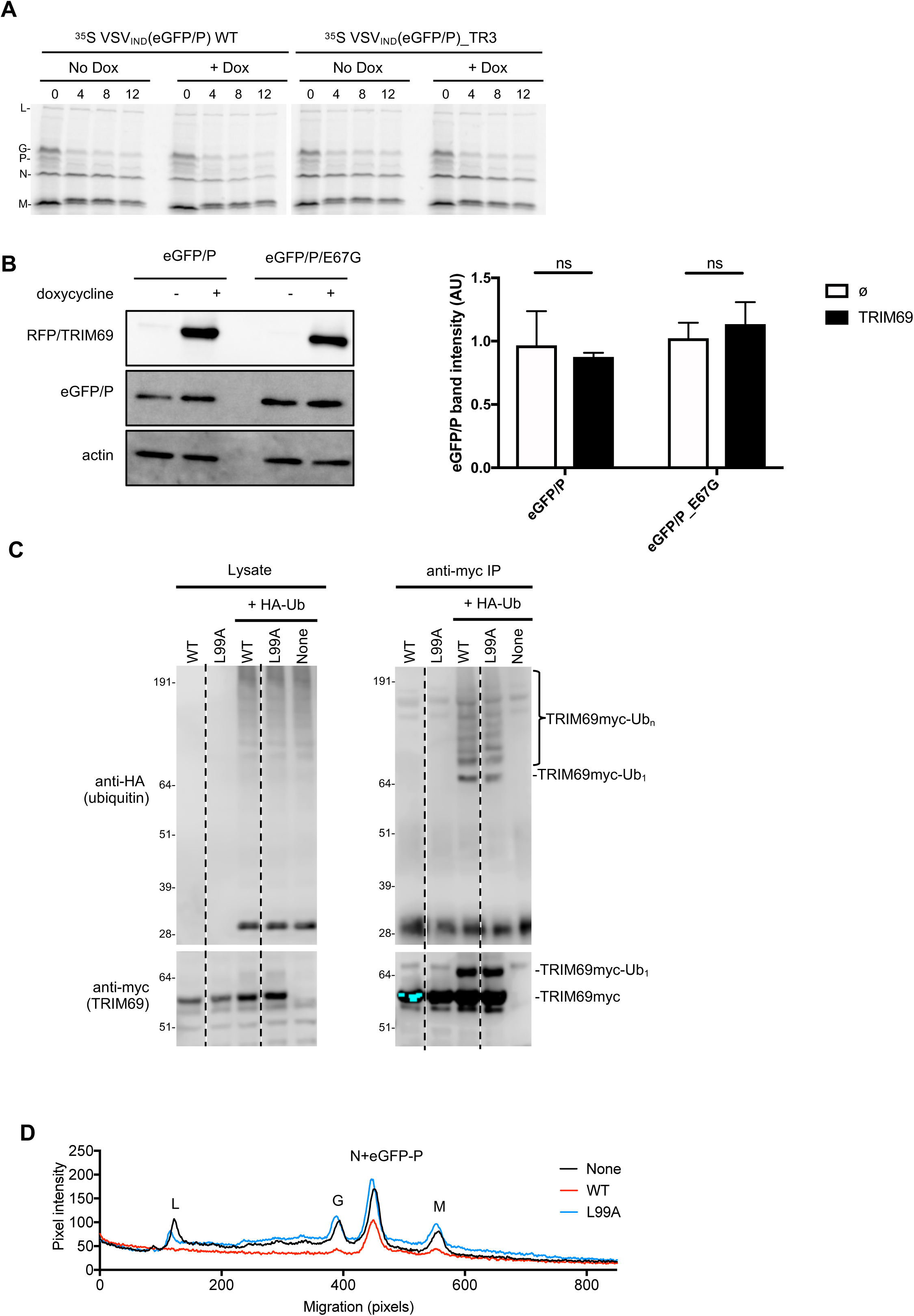
(A) HT1080-TagRFP/TRIM69 cells were seeded and simultaneously treated or not with doxycycline. Sixteen hours later, cells were treated with cycloheximide and infected with ^35^S radio-labelled VSV_IND_(eGFP) or VSV_IND_(eGFP/P)_TR3(E67K) at MOI 10. Cells were harvested at 0, 4, 8 and 12 h.p.i. and proteins were analyzed by SDS-PAGE. (B) Western blot analysis of eGFP/P protein levels in transfected HT1080-TagRFP/TRIM69 cells that were treated or not with doxycycline. Left panel represents a representative blot, right panel shows quantification of eGFP/P band intensities (mean ± sd, n=4) (C) TRIM69 autoubiquitination in 293T cells co-transfected with plasmids expressing TRIM69/myc and HA-ubiquitin. Lysates (left) and anti-myc immunoprecipitates (right) were probed with anti-myc and anti-HA tag antibodies. (D) Quantification of pixel intensities versus migration for the leftmost three lanes in Fig. 6C, extracted using ImageJ

### SI Materials and Methods

#### Cells

HT1080 cells were obtained from ATCC and maintained in Dulbecco’s Modified Eagles Medium (DMEM) (ThermoFisher Cat# 11995065) with 10% fetal calf serum (FCS) (Sigma, Cat# F8067-500ML) and gentamicin (ThermoFisher Cat# 15710064). An IFN*α* (PBL Assay Science Cat# 11200-2) sensitive HT1080 single cell clone was derived by limiting dilution. BHK21 cells were obtained from ATCC maintained in Eagle’s Minimum Essential Medium (EMEM) (ATCC Cat# 30-2003) supplemented with 10% FCS (Sigma, Cat# F8067-500ML) and gentamicin (ThermoFisher Cat# 15710064). Cells were not tested for mycoplasma contamination, but were periodically tested for retrovirus contamination using a PCR-based reverse transcriptase assay (1, 2)

BSRT7 cells (a kind gift from K. Conzelmann) (3) and African green monkey kidney Vero cells (ATCC, CCL-81) were maintained in Dulbecco’s modified Eagle’s medium (DMEM; Corning Inc., 10-013-CV) containing 10 % fetal bovine serum (FBS; Tissue Culture Biologicals, TCB 101) at 37 °C and 5 % CO_2_.

#### Plasmid construction

The pLKO-derived doxycycline-inducible lentiviral expression vector was used as previously described (4). pLKO TagRFP or mScarlet N-terminally tagged human TRIM69, various species and mutants were cloned by overlapping PCR using SfiI restriction sides. pLKO myc-TRIM69 or mutant constructs were cloned using a forward primer containing a myc tag and SfiI restriction sites. Plasmids expressing WT or mutant TRIM69-3xmyc were generated in the pCR3.1 expression vector containing a multiple cloning site EcoRI-XhoI-NotI followed by in frame 3xmyc tag using EcoRI and XhoI restriction sites. Plasmids expressing WT or mutant Cherry-TRIM69 were generated in the pCR3.1 Cherry EcoRI-XhoI-NotI background using EcoRI and XhoI restriction sites.

pCAGGS-eGFP/P_TR3 (E67G) was derived from pCAGGS-eGFP/P (5) and was made by site-directed mutagenesis using the Q5® High fidelity DNA polymerase (New England Biolabs, M0491) and the primers ATTGAAGACAATCAAGGCTTGTATG and TTGATTGTCTTCAATacCTGGTTCAGATTCTGTGTCAGAAT.

Sequences encoding VSV-P WT and mutants E67G and E69K amino acid 32 to 107 were amplified from pCAGGS-eGFP/P plasmids and inserted in frame with the Glutathione-S-transferase (GST) in the pCAGGS-GST expression plasmid using the following oligonucleotide primers and restriction enzymes (5’ EcoRI GAGGAGGAATTCGCTGAAAAGTCCAATTATGAGTTG, 3’ XhoI CTCCTCCTCGAGCTAGTCCGAAGTAAATACAACATCCAC).

The plasmids used to produce Sendai virus were kindly provided by Benhur Lee. The provided rSeV clone contained eGFP and mutations in the F and M genes to allow trypsin-independent growth. A Hh-Rbz sequence was present between the T7 promoter and the start of the viral antigenome to enhance the rescue efficiency (6). The GFP gene which had been positioned between the N and P genes via duplication of the N-P intergenic region was replaced with nLuc. Briefly, the N and P region were amplified with Pre-SbfI-for (5’ TGACCATGATTACGCCAAGCTTAA 3’) to nLuc_SeV_rev (5’ GAAATCTTCGAGTGTGAAGACCATGCGGTAAGTGTAGCCGAAGCCGTG 3’) and nLuc-SeV_for (5’ CTGTGCGAACGCATTCTGGCGTAATGAgatagGAGGAATCTAGGATCA 3’) to Post-SmaI-6860-rev (5’ GATGGTAGATTGGGTCTCTCTGTG3’), respectively. The nLuc gene (Promega) was amplified with SeV-nLuc_for (5’ CACGGCTTCGGCTACACTTACCGCATGGTCTTCACACTCGAAGATTTC 3’) and SeV-nLuc_rev (5’ TGATCCTAGATTCCTCCTATCTCATTACGCCAGAATGCGTTCGCACAG 3’). After overlap extension PCR using the most outer primers, the fragment was cloned into the with Sbf1 and Sma1 linearized rSeV_GFP construct and combined using Gibson assembly. pVSV_NJ_(+)-eGFP was construct as previously described for plasmid pVSV1(+) encoding VSV_IND_ genomic RNA (7). Briefly, pVSV_NJ_(+)-eGFP was assembled from plasmid made by reverse transcription-PCR of each of the VSV_NJ_ genes of the Ogden strain and intergenic junctions by standard cloning techniques. These clones were assembled into a full-length cDNA and inserted between the bacteriophage T7 promoter and a cDNA copy of the self-cleaving ribozyme from the antigenomic strand of HDV. The eGFP gene was inserted in first position (in an addition transcription unit before N) as described for VSV_IND_ (8).

#### Viruses

Plasmids encoding the full length VSV_IND_ genome (pVSV-FL) as well as individual VSV_IND_ genes N, P, L, and G were purchased from Kerafast (VSV-FL+[2] VSV Plasmid Expression Vector System, EH1002) or were generated as described previously (7). VSV_IND_ viruses were generated by infecting 293T cells with T7-expressing vaccinia (vTF7-3) at a MOI of 5, followed by transfection with pVSV plasmids and plasmids encoding VSV-N, P, L, and G under the control of a T7 promoter. Supernatants were harvested 48 h.p.t., filtered (0.2 μm) to remove the bulk of the vaccinia virus and plaque purified on 293T cells. Plaque purified virus was expanded on 293T cells and cell culture supernatant was harvested, passed through a 0.2 μm filter and frozen in aliquots. Virus titers (PFU/mL) were determined by plaque formation using HT1080 or BHK21 cells. VSV encoding nanoluciferase (nLuc) were generated by inserting the nLuc encoding sequences (from pNL1.1, Promega) into the pVSV plasmid between the envelope and L genes, along with appropriate VSV regulatory sequences. VSV derivatives encoding mNeonGreen were generated by fusing the mNeonGreen encoding sequence to the N terminus of P.

VSV_IND_(eGFP) (8), VSV_IND_(eGFP/P) (5) and RabV(eGFP-ΔG) (9) were described previously. VSV_NJ_(eGFP) was rescued from pVSV_NJ_(+)_eGFP following a previously described protocol (7) Rhabdovirus stocks were grown on BSR-T7 cells, BHK cells or Vero cells, and titered by plaque assay on BSR-T7 or HT1080 cells. Briefly, cells were seeded in DMEM - 10 % FBS and infected one day later for 1 h with viruses at a multiplicity of infection (MOI) of 0.01. Virus suspensions were replaced by DMEM - 2 % FBS and cell supernatants were harvested when 95 % of the cells were infected and ready to detach (about 24 h for VSV_IND_). For RabV(eGFP-ΔG)-G_VSVind_, cells were transfected 8 hours prior infection with plasmid expressing VSV_IND_-G using Lipofectamine^TM^ 2000 (InvitrogenTM Cat# 11668-019). For concentrated sucrose cushion-purified virus stocks, infected cell supernatant was concentrated through a 15 % sucrose cushion in NTE (10 mM Tris-HCl pH 7.4, 100 mM NaCl, 1 mM EDTA) at 110,000 x g for 2 h at 4°C. Pellets were resuspended overnight at 4 °C in NTE.

Rescue of replication-competent Sendai virus from transfected plasmids was done as previous described (6) with transfection into 293T cells using Lipofectamine LTX (Invitrogen Cat# 15338100) according to manufacturer’s recommendations. Virus titers (PFU/mL) were determined by plaque formation using HT1080 target cells.

#### siRNA based ISG screen

The 400 most IFN*α* inducible ISGs were chosen using a compilation of microarray data from 293T, HT1080, CEM, Jurkat, MT2, MT4, C8166, Hut R5, H9, Sup T1, U937, THP-1, K562, HL60 and KG1a cells, and are listed in SI appendix Table S1. All ISG screens were conducted in a 96-well format. 3×10^3^ HT1080 cells were plated and transfected with siRNA SMARTpools (Dharmacon) using RNAimax (Invitrogen Cat# 13778150) the following day and treated with 10 U/ml of IFN*α* (PBL Assay Science Cat# 11200-2) 8 h.p.t.. The cells were infected with 30 PFU per well of VSV nLuc the next day. At 20 h.p.i., cells were washed 3 times in 1x PBS, lysed in passive lysis buffer (Promega Cat# E1941) and luciferase was measured using the Nano-Glo Luciferase Assay System (Promega Cat# N1130) and Modulus II Microplate Multimode Reader (Turner BioSystems).

#### siRNA experiments

3×10^3^ HT1080 cells were plated into a 96-well plate and transfected with siRNA smart pools or most efficient individual siRNA (Dharmacon see Table 1) and treated with increasing concentrations of IFN*α* or a fixed dose of 10 U/ml of IFN*α* (PBL Assay Science Cat# 11200-2) 8 h.p.t. and harvested as described above.

#### Inducible expression of TRIM69

Inducible HT1080 cells were generated by transduction with a LKO-derived vector as described (4) followed by selection with 1.25 μg/mL puromycin (Sigma-Aldrich Cat# P8833-100MG). Expression was induced in pLKO transduced cell lines through an overnight treatment with 0.5 μg/mL doxycycline hyclate (Sigma-Aldrich Cat# 324385) prior to viral challenge.

#### Deconvolution and Structured illumination Microscopy (SIM)

3×10^4^ HT1080 cells were plated onto gelatin coated 8-chambered #1.5 borosilicate glass bottomed slides (LabTek Cat#155409) and TagRFP/TRIM69 or mScarlet/TRIM69 expression was induced by overnight treatment with 0.5 μg/mL doxycycline hyclate (Sigma-Aldrich). Cells were infected with mNeonGreen/P or eGFP/P VSV at an MOI of 3 and fixed 4 h.p.i. using 4% formaldehyde (Cat# Sigma P6148-1KG) and imaged by deconvolution microscopy (DeltaVision OMX SR imaging system). All images were generated by maximum intensity projection using the Z project function in ImageJ (Version 2.0.0-rc-59/1.51w).

#### Confocal Microscopy

HT1080 TagRFP/TRIM69 cells were seeded on a 1.5 mm coverslip (Warner Instruments Cat# CS-12R15) in a 24-well plate and simultaneously treated or not with 0.5 μg/mL doxycycline. Sixteen hours later, cells were transfected or not with pCAGGS-eGFP/P and pCAGGS-eGFP/P(E67G) plasmids with Lipofectamine^TM^ 2000 (Invitrogen^TM^ Cat# 11668-019) following manufacturer’s protocol. Twenty-four hours post-transfection, cells were washed with 500 μL Dulbeco’s Phosphate Buffered Saline liquid (DPBS; Sigma Cat# 59300C), fixed for 15 min at room temperature with 250 μL DPBS containing 2 % paraformaldehyde, washed twice with 1 mL DPBS containing 10 mM glycine (to quench residual paraformaldehyde) stained for 15 min with DAPI diluted in DPBS containing 0.5 % bovine serum albumin, washed twice with 1 mL DPBS containing 0.05 % Tween-20 and once with 1 mL H2O. Coverslips were mounted onto slides with 4 μL ProlongGold (Invitrogen Cat# P36930). Confocal images were acquired on a Nikon T1 inverted microscope equipped with a Yokogawa CSU-W1 scanhead, Toptica laser launch and an Andor Zyla 4.2 plus sCMOS camera using a plan apo 100x lambda/1.45 DIC oil objective. The acquisition software was NIS Elements AR 5.02. The emitted light from eGFP and TagRFP fluorophores was collected using a Semrock multiband pass dichroic filter (Di01-t 405/488/561/647) and Chroma 525/36 and 605/52 band pass emitter, respectively.

#### Single molecule FISH

smFISH probes against both the plus and minus strands of VSV N and P and were designed using the Stellaris Probe Designer Version 2.0 (SI Appendix Table S2) (https://www.biosearchtech.com/stellaris-designer). For each RNA, 29 – 41, oligonucleotide probes were synthesized by IDT to contain a 5’ Amino Modifier (C6). The 5’Amino modified probes for each RNA were resuspended to 1.25 μg/mL, pooled, and purified by three chloroform extractions followed by ethanol precipitation. Then, 50 μg of the pooled probes were labeled with ester-modified Alexa-488 or Alex-549 using the Alexa Fluor 488 Oligonucleotide Amine Labeling Kit (Thermo Fisher Cat# A20191). After labeling the pooled probes were ethanol precipitated, resuspended in RNase-free water and purified via the Oligo Clean & Concentrator Kit from Zymo Research (Cat# D4060). The pool probes were eluted in RNase-free TE pH 8.0 (Ambion Cat# AM9849) and adjusted to a final concentration of 12.5 μM. For FISH, 3×10^4^ HT0180-myc/TRIM69 cells were seeded onto gelatin coated 8-chambered #1.5 borosilicate glass bottomed slides (LabTek Cat# 155409). Doxycycline treated or untreated cells were pretreated for 30 min with 100 μg/mL of cycloheximide (Sigma-Aldrich Cat# C4859) and infected at an approximate MOI of 20 with VSV(mNeonGreen/P) virus. 2h and 45 min post-infection the cells were washed with PBS (Ambion Cat# AM9624) and fixed with 4% formaldehyde (ThermoFisher Cat# 28908) in PBS for 30 min at RT. Following permeabilization with 70% ethanol for 2h at RT the cells were briefly washed with Stellaris RNA FISH Wash Buffer A (Cat# SMF-WA1-60); for 5 min at RT. The cells were probed for N or P plus or minus strand RNA with the 0.125 μM Alexa-488 or Alex-549 labeled probes in Stellaris RNA FISH Hybridization Buffer (Cat# SMF-HB1-10) for 16-18h at 37°C. The cells were then washed two times for 30 min at 37°C in Stellaris RNA FISH Wash Buffer A (Cat# SMF-WA1-60); the second wash contained Hoechst at 1 μg/mL. After a 5 min wash with Stellaris RNA FISH Wash Buffer B (Cat# SMF-WB1-20) cells were rinsed three times with PBS and imaged by deconvolution microscopy (DeltaVision OMX SR imaging system). All images were generated by maximum intensity projection using the Z project function in ImageJ (Version 2.0.0-rc-59/1.51w). RNA spots were quantified using StarSearch developed by the Raj laboratory (http://rajlab.seas.upenn.edu/StarSearch/launch.html).

#### VSV replication assays

1×10^4^ HT1080 cells were plated in a 96-well plate format and TRIM69, Mx1, CD68 or RFP expression was induced by overnight treatment with 0.5 μg/mL doxycycline hyclate (Sigma-Aldrich). The cells were infected with 30 PFU per well of VSV(nLuc) the next day. At 20 h.p.i. or indicated timepoints, supernatant was collected and cells were lysed in passive lysis buffer (Promega Cat# E1941) and luciferase was measured using the Nano-Glo Luciferase Assay System (Promega Cat# N1130) and Modulus II Microplate Multimode Reader (Turner BioSystems). Or supernatant containing virions were titered on BHK21 cells under a methyl cellulose overlay.

To compare the growth of the TR viruses, 1.2×10^6^ Vero cells were infected with the different viruses at MOI 0.05. Aliquots of the supernatant were harvested at 8, 12, 16, 20 and 24 h.p.i. and titered by cytometry on BSR-T7 cells. Titers are expressed in infectious unit per mL: the number of virions leading to detectable expression of eGFP in BSR-T7 cells, per mL. For short term (single cycle) infection assays, HT1080-TagRFP/TRIM69 cells were seeded and simultaneously treated or not with 0.5 μg/mL doxycycline. Sixteen hours later, cells were infected with VSV_IND_(eGFP), VSV_NJ_(eGFP) or RabV(eGFP-ΔG)-G_VSVind_ at MOI = 1 for 1 h. Six hours after infection, TagRFP/TRIM69 and eGFP expression levels were monitored by epifluorescence microscopy.

#### Selection of TRIM69 resistant (TR) viruses

TRIM69-resistant VSV_IND_(eGFP) and VSV_IND_(eGFP/P) were selected by plaque assay on HT1080-TagRFP/TRIM69 cells. Cells were seeded and simultaneously treated with 0.5 μg/μL doxycycline (Sigma Cat# D9891). Sixteen hours later, cells were infected for 1h with 1:10 dilutions of the viral stocks and overlaid with medium containing 0.25 % agarose. Plaque were picked and amplified once on HT1080 expressing TagRFP/TRIM69 and then on BSR-T7 cells.

#### Western blotting

For Fig 2A, 5B and SI Appendix Fig. S1A, C, S9C, cells were lysed in LDS sample buffer (Invitrogen Cat# NP0008) and proteins were separated by electrophoresis on NuPage 4-12% Bis-Tris gels (Invitrogen Cat# NP0323BOX) and blotted onto nitrocellulose membranes (GE Healthcare Cat# 10600003). Membranes were incubated with rabbit anti-HSP90 (Proteintech Cat# 13171-1-AP), mouse anti-VSV-M (Kerafast Cat# EB0011), mouse anti-RFP (Abcam Cat# ab125244), mouse anti-myc (Biolegend Cat# 904401), rabbit anti-TRIM69 (Proteintech Cat# 12951-1-AP) or rabbit anti-CD68 (Proteintech Cat# 25747-1-AP). Thereafter, membranes were incubated with goat anti-rabbit IRDye 800CW and goat anti-mouse IRDye 680RD (LI-COR Biosciences Cat#926-32211 and 926-32220 respectively), and scanned using a LI-COR Odyssey infrared imaging system. Alternatively, membranes were incubated with appropriate horseradish peroxidase (HRP) conjugated secondary antibodies (Jackson ImmunoResearch Goat Anti Mouse Cat# 15-035-174 Goat Anti Rabbit Cat# 111-035-144), and visualized using SuperSignal West Femto Chemiluminescent solution (ThermoFisher Cat# PI34095) and a C-DiGit Western Blot Scanner (LiCor).

For SI Appendix Fig. S9B, cells were lysed in 50 μL of 20 mM Tris-HCl pH 8, 150 mM NaCl, 0.6 % NP-40, 2 mM EDTA and 1X protease inhibitor cocktail (cOmpleteTM, Roche Cat# 4693116001). Soluble proteins were separated on 10 % acrylamide gels, transferred onto nitrocellulose membranes and incubated with either mouse anti-RFP (Abcam Cat# ab125244), rabbit anti-GFP (Abcam Cat# ab6556) or mouse anti-actin (Sigma Cat# A5316) antibodies followed by incubation with HRP conjugated anti-mouse (Invitrogen Cat# 31430) or anti-rabbit antibodies (Sigma Cat# A0545). HRP activity was visualized using Pierce ECL Western kit (Thermo Scientific Cat# 32209) and imaged with Amersham Imager 600 (GE Healthcare). Protein bands intensities were quantified using ImageJ software.

#### Glutathione-S-transferase coprecipitation assay

Approximately four million 293T cells were transfected via Polyethylenimine (Polysciences Cat# 23966) with 5 μg or GST, GST/P(32–107) (WT, E69K or E67G) and 5 μg Cherry, Cherry/TRIM69 WT or L99A. Two days post transfection the cells were lysed on ice for 10 min in 1 mL 50 mM Tris pH 7.4, 150 mM NaCl, 1% digitonin. Lysates were cleared by centrifugation and incubated with 25 μL Glutathione Sepharose 4B beads (GE Healthcare Cat# 17075601) for 3h at 4°C. The beads were washed 3 times in 1 mL lysis buffer and eluted by boiling in 50 μL 1X SDS sample buffer. The eluted proteins were separated on NuPage 4-12% Bis-Tris Protein Gel (Thermo Fisher Cat# NP0323BOX) and transferred onto nitrocellulose (GE Healthcare Cat# 10600003) for probing with rabbit anti-GST (Abcam Cat# 19256) and mouse anti-RFP (Abcam Cat#125244) follow by goat anti-rabbit-IRDye 680 (LiCor Cat# 926-68071) and goat anti mouse-IRDye 800 (LiCor Cat# 926-32210).

#### Measurement of TRIM69 multimerization

Inducible HT1080 cells (3×10^6^) expressing wild type or mutant TagRFP/TRIM69 were collected in 1X PBS and crosslinked by treatment with 0.2 mM EGS (CovaChem Cat# 13308-100), a membrane-permeable crosslinker. After 30 min of incubation at room temperature, cells were lysed in LDS sample buffer (Invitrogen), proteins were separated by electrophoresis on NuPage 4-12% Bis-Tris gels (Invitrogen Cat# NP0323BOX) and blotted onto nitrocellulose membranes (GE Healthcare Cat# 10600003). Blots were probed with rabbit anti-HSP90 (Proteintech) and mouse anti-RFP (Abcam Cat# 125244). Thereafter, membranes were incubated with goat anti-rabbit IRDye 800CW and goat anti-mouse IRDye 680RD (LI-COR Biosciences Cat#926-32211 and 926-32220 respectively), and scanned using a LI-COR Odyssey infrared imaging system.

#### TRIM69 autoubiquitination

7×10^5^ 293T cells were transfected via Polyethylenimine (Polysciences Cat# 23966) with 1 μg of pCR3.1 TRIM69-3xmyc and 500 ng of pHA-ubiquitin. At 36 h.p.t., cells were thoroughly lysed at room temperature in detergent-rich RIPA buffer (50mM Tris pH 7.4, 150 mM NaCl, 1mM EDTA, 1.0% glycerol, 0.5% SDS, supplemented with protease inhibitor (Roche Cat# 04693159001) and 5 mM N-ethylmaleimide (Sigma Cat# 04259-5G), to inhibit de-ubiquitination), sonicated and cleared of cellular debris by microcentrifugation. Lysates were transferred into fresh Eppendorf tubes and diluted 5-fold in the same buffer containing NP-40 rather than SDS, to adjust the concentration of SDS to 0.1% and NP-40 to 1.0%. Lysates were incubated with 30 μL Dynabeads (Invitrogen Cat# 10001D) for 2h at 4°C. The beads were washed 3 times in 1 mL lysis buffer and eluted by boiling in 50 μL 1X SDS sample buffer. The eluted proteins were separated on NuPage 4-12% Bis-Tris Protein Gel (Thermo Fisher Cat# NP0323BOX) and transferred onto nitrocellulose (GE Healthcare Cat# 10600003) for probing with mouse anti-myc (Biolegend Cat# 904401) and rabbit anti-HA (Proteintech Cat# 51064-2-AP) followed by goat anti-rabbit IRDye 800CW and goat anti-mouse IRDye 680RD (LI-COR Biosciences Cat# 926-32211 and 926-32220 respectively), and scanned using a LI-COR Odyssey infrared imaging system.

#### Radio-labelling and analysis of virion proteins

For production of virions containing radio-labelled proteins, BSR-T7 cells were seeded in a 150mm dish in DMEM - 10 % FBS. The next day, cells were incubated with 4 mL methionine-free, cysteine-free DMEM (Corning Inc. Cat# 17-204-Cl) for 30 min and infected for 1h at MOI 3 in 4 mL methionine-free, cysteine-free DMEM. Virus solution was then replaced with 12 mL methionine-free, cysteine-free DMEM containing 120 μL of EXPRE35S35S Protein Labeling Mix (PerkinElmer Cat# NEG072007MC). Twenty hours later, cell supernatants were harvested, sucrose cushion-purified and viruses were tittered by plaque assay.

To monitor virion protein decay, TagRFP/TRIM69 cells were seeded in a 12-well plate and simultaneously treated or not with 0.5 μg/mL doxycycline. Sixteen hours later, cells were treated with 100 μg/mL cycloheximide and infected for 1 h with 35S radio-labelled viruses at MOI 10. Cells were washed twice with DMEM – 2% FBS and incubated in 0.5 mL DMEM – 2 % FBS containing 100 μg/mL cycloheximide. At 0, 4, 8 and 12 h.p.i., cells were harvested, washed in DPBS and lysed with 20 μL Rose lysis buffer (50 mM Tris-HCL pH 7.4, 5 mM EDTA, 150 mM NaCl, 1 % NP-40, 1 X protease inhibitor cocktail (cOmpleteTM, Roche Cat# 4693116001)). Proteins were analyzed by SDS-PAGE, gels were fixed (in 30% methanol and 10% acetic acid), dried, exposed overnight to a phosphor screen, and the radiolabeled proteins were visualized using a Typhoon FLA 9500 scanner. Protein band intensities were quantified with ImageJ software.

#### Radio-labelling and analysis of primary transcripts

HT1080-TagRFP/TRIM69 and HT1080TagRFP/TRIM69(L99A) cells were seeded in 6-well plates and simultaneously treated or not with 0.5 μg/mL doxycycline. Sixteen hours later, cells were incubated in phosphate-free DMEM (Gibco Cat# 11971-025) for 30 min followed by a 30 min incubation in phosphate-free DMEM containing or not 0.5 μg/mL doxycycline, 10 μg/mL actinomycin-D (Sigma Cat# A5156) and 100 μg/mL cycloheximide (VWR Cat# 94271). Cells were then infected for 30 min with sucrose cushion-purified virus at MOI 100. Virus solutions were replaced by 1 mL phosphate-free DMEM containing or not 0.5 μg/mL doxycycline, 10 μg/mL actinomycin-D and 100 μg/mL cycloheximide, and 10 μL of Phosphorus-32 Radionuclide (PerkinElmer Cat# NEX053H005MC) were added dropwise. Five hours post infection, RNA were extracted using Trizol reagent (Invitrogen Cat# 15596018) following manufacturer’s protocol. RNA were boiled at 100 °C for 1 min, incubated on ice for 2 min, mixed with a 1.33 X loading buffer (33.3 mM citrate pH 3, 8 M urea, 20 % sucrose, 0.001 % bromophenol blue) and analyzed on a 25 mM citrate pH 3, 1.75 % agarose, 6 M urea gel ran for 18h at 4 °C and 180 V. Gels were fixed (in 30% methanol and 10% acetic acid), dried, exposed overnight to a phosphor screen (GE Healthcare), and the radiolabeled RNA products were visualized using a Typhoon FLA 9500 scanner (GE Healthcare).

---

**Table S1.**
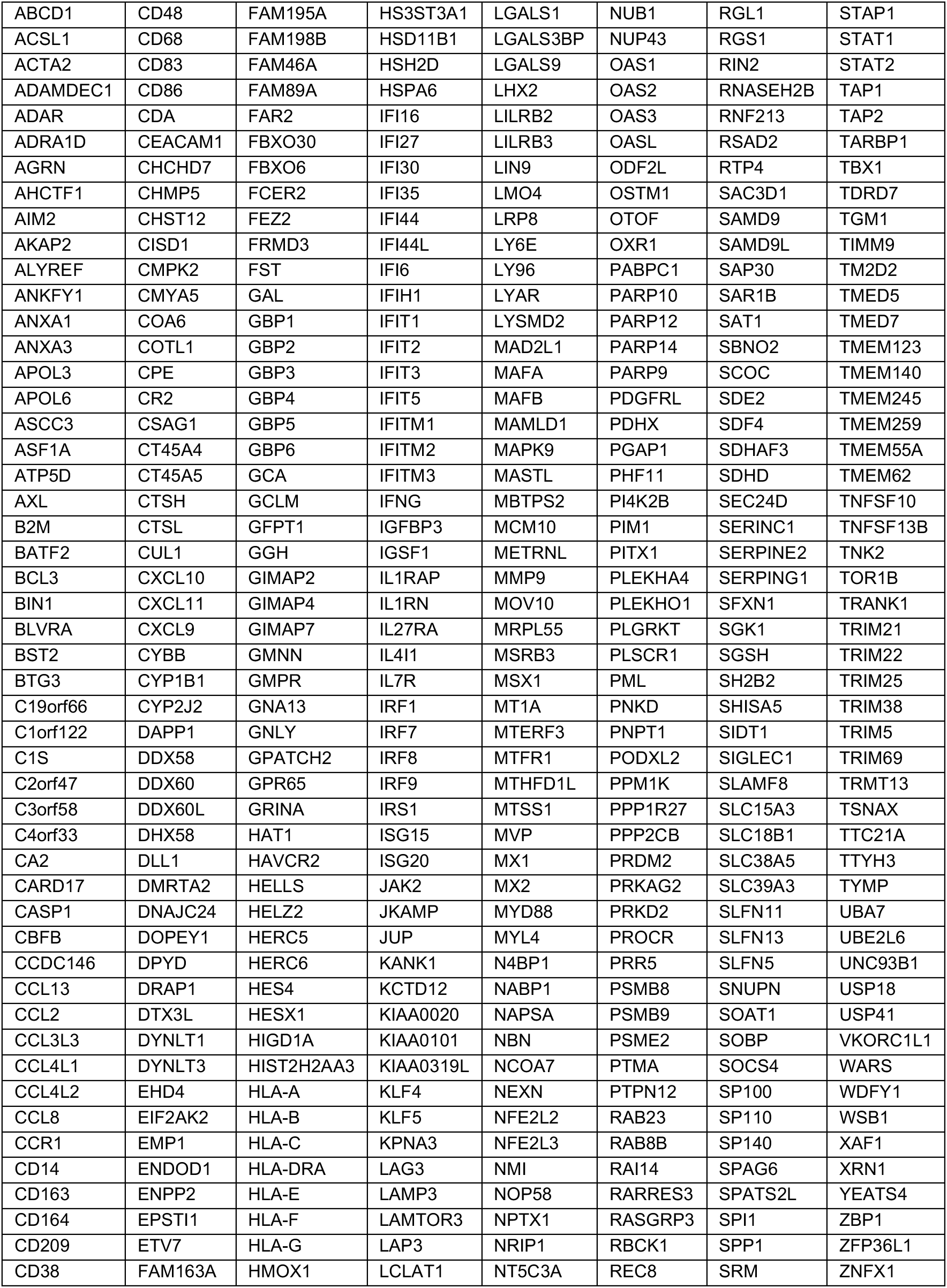
ISG SMARTpools included in siRNA screen.

**Table S2.**
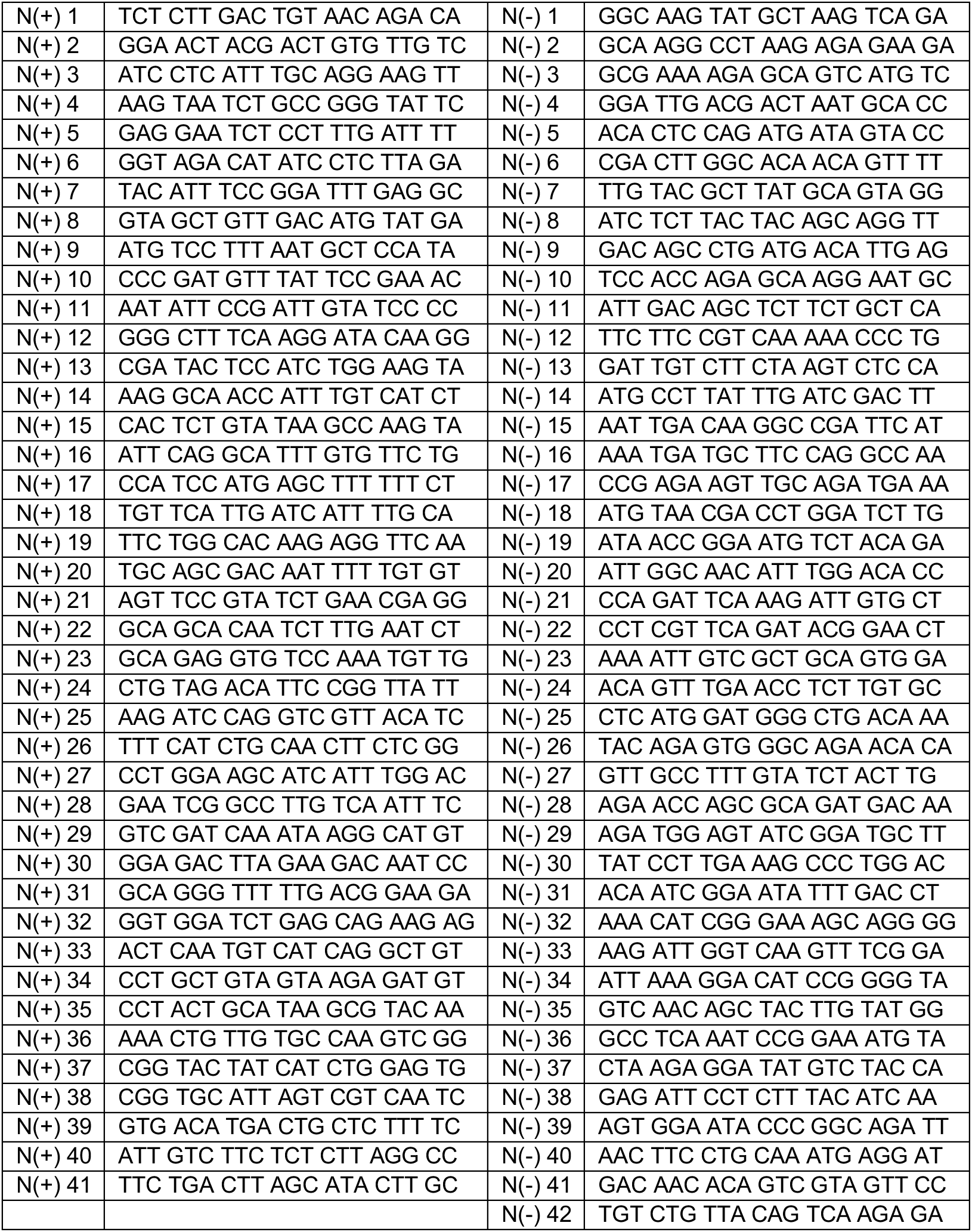
FISH probes targeting VSV_IND_ N positive and negative RNA strands.

